# scBSP: A fast and accurate tool for identifying spatially variable features from high-resolution spatial omics data

**DOI:** 10.1101/2025.02.02.636138

**Authors:** Jinpu Li, Mauminah Raina, Yiqing Wang, Chunhui Xu, Li Su, Qi Guo, Ricardo Melo Ferreira, Michael T Eadon, Qin Ma, Juexin Wang, Dong Xu

**Author notes:** These authors contributed equally (Jinpu Li, Mauminah Raina). Corresponding Authors Qin Ma, Juexin Wang, Dong Xu.

## Abstract

Emerging spatial omics technologies empower comprehensive exploration of biological systems from multi-omics perspectives in their native tissue location in two and three-dimensional space. However, sparse sequencing capacity and growing spatial resolution in spatial omics present significant computational challenges in identifying biologically meaningful molecules that exhibit variable spatial distributions across different omics. We introduce scBSP, an open-source, versatile, and user-friendly package for identifying spatially variable features in high-resolution spatial omics data. scBSP leverages sparse matrix operation to significantly increase computational efficiency in both computational time and memory usage. In diverse spatial sequencing data and simulations, scBSP consistently and rapidly identifies spatially variable genes and spatially variable peaks across various sequencing techniques and spatial resolutions, handling two- and three-dimensional data with up to millions of cells. It can process high-definition spatial transcriptomics data for 19,950 genes across 181,367 spots within 10 seconds on a typical desktop computer, making it the fastest tool available for handling such high-resolution, sparse spatial omics data while maintaining high accuracy. In a case study of kidney disease using 10x Xenium data, scBSP identified spatially variable genes representative of critical pathological mechanisms associated with histology.

## INTRODUCTION

Identifying spatially variable genes (SVGs) is one of the fundamental tasks in spatially resolved transcriptomics (SRT) studies for understanding tissue organization and cell regulation^1^. Beyond SVGs on the mRNA level, emerging spatial omics techniques provide a more comprehensive view of measuring diverse molecular features as spatially variable features (SVFs) across different levels of multi-omics, which includes genomics, epigenomics, transcriptomics, proteomics, lipidomics, and metabolomics^2–4^. By describing biological variances in the space from different angles of biological processes, SVFs play a crucial role in comprehending tissue structures and functions ^5–11^. However, these fast-accumulating spatial omics data in high-dimensionality and scalability raise considerable computational gaps in identifying biologically meaningful SVFs within the tissue context in multiple omics.

Diverse characteristics of multiple modalities are the first challenge in spatial omics analysis. It is typically difficult to have one computational approach to uniformly model modalities with diverse data qualities and distribution characteristics, such as modeling spatially profiled gene and protein expression, and chromatin opening structure. The second challenge comes from growing spatial resolution. Advancements in high-resolution SRT, including Slide-seq ^12^, high-definition spatial transcriptomics (HDST) ^13^, Visium HD, 10x Xenium, NanoString CosMx, MERSCOPE, and Stereo-seq (15), have propelled transcriptome-wide profiling to a single-cell or subcellular resolution with a more significantly increased number of spots or cells ^15,16^.

Meanwhile, spatial epigenomics, such as Spatial ATAC-seq (Assay for Transposase-Accessible Chromatin using sequencing) and spatial-CUT&Tag, bring at least one order of magnitude of feature numbers as ATAC-seq peaks rather than genes, challenging the capacities of existing computational methods. Beyond the early years of two-dimensional (2D) slices, registration and modeling of three-dimensional (3D) contexts is the third challenge in spatial omics research. 3D techniques like STARmap ^17^ have demonstrated substantial advantages in a more comprehensive and faithful representation of intact organ structures and functions, enhancing accurate quantitative interpretation ^17–23^. However, the shift from 2D to 3D space brings new challenges in data analysis, including the significantly growing number of spatial locations and a high proportion of zero values with limited sequencing depth ^24–27^. In addition, appropriately handling the sparsity in these high-resolution, high-dimensional multi-omics data is still an everlasting challenge in methods development.

Due to these computational challenges, although numerous methods and tools have been developed to identify SVGs from SRT, few can be directly applied to detecting SVFs in other omics modalities, especially in spatial epigenomics research, where peaks in spatial ATAC-seq can even reach millions^28^. Tools such as nnSVG ^29^ and SOMDE ^30^ were optimized to scale linearly with the increasing number of spatial locations, but their computational efficiency is still insufficient for handling spatial omics data encompassing hundreds of thousands of spatial locations with hundreds of thousands of features. Specially designed methods such as SpaLDVAE also struggle to handle spatial ATAC-seq data on an even larger scale. In addition, the violation of model assumptions and computational complexity hinders the application of current methods in spatial omics, especially on complex 3D omics data. For example, due to inconsistencies in inter-plane and within-plane spatial resolution, identifying SVGs from 3D SRT has limited accuracy based on existing tools like SPARK-X ^25^ and spaGCN ^31^.

Here, we present **scBSP** (single-cell big-small patch), a significantly enhanced implementation of our previous method BSP ^32^, to address computational challenges in identifying SVFs from high-resolution two/three-dimensional spatial omics data. scBSP selects a set of neighboring spots within a certain distance to capture the regional means and filters the SVFs based on the rate of changes in the variance of these local means with different granularities. Incorporating sparse matrix techniques and approximate nearest neighbor search^33^, scBSP significantly reduces computational time and memory consumption compared to the original implementation of BSP and other existing approaches, reducing the processing time to seconds for high-resolution SRT HDST data. In diverse case studies with multiple sequencing platforms, scBSP accurately identifies biologically meaningful SVFs within seconds to minutes on a personal laptop, including handling spatial ATAC-seq data with hundreds of thousands of peaks without sacrificing efficiency. Notably, scBSP stands out for its robustness, as it does not assume any specific distribution of gene expression levels or spatial patterns of spots, and it does not have model parameters to tune or train. This characteristic ensures its adaptability and robustness to various spatial omics sequencing techniques, ensuring consistent results without the need to adapt the model and effectiveness in addressing the zero-inflation issue commonly encountered in high-resolution data analyses^24,25^.

## RESULTS

### scBSP accelerates computational efficiency by a thousand-fold on high-resolution spatial omics data, while maintaining comparable performance as BSP

scBSP is an open-source software for identifying SVFs in dimension-agnostic and technology-agnostic spatial omics data. Its technical details are in the “Methods” and method schematic shown in **Figure 1A**. We systematically evaluated and compared the computational efficiency and model accuracy of scBSP with seven competitive methods, including Moran’s I, spatialDE, SPARK, SPARK-X, nnSVG, SOMDE, and BSP, for detecting SVGs on simulated SRT data (simulation details are provided in “Methods”). Two simulation scenarios were designed to evaluate computational efficiency for spatial omics data at both spot-level and cellular-level resolution. The model accuracy was assessed using statistical power on the simulated benchmarks. To be compatible with the high memory usage of competitive methods, all methods were executed on a workstation with a 2.00GHz AMD EPYC 7713 64-Core Processor.

**Figure 1:**
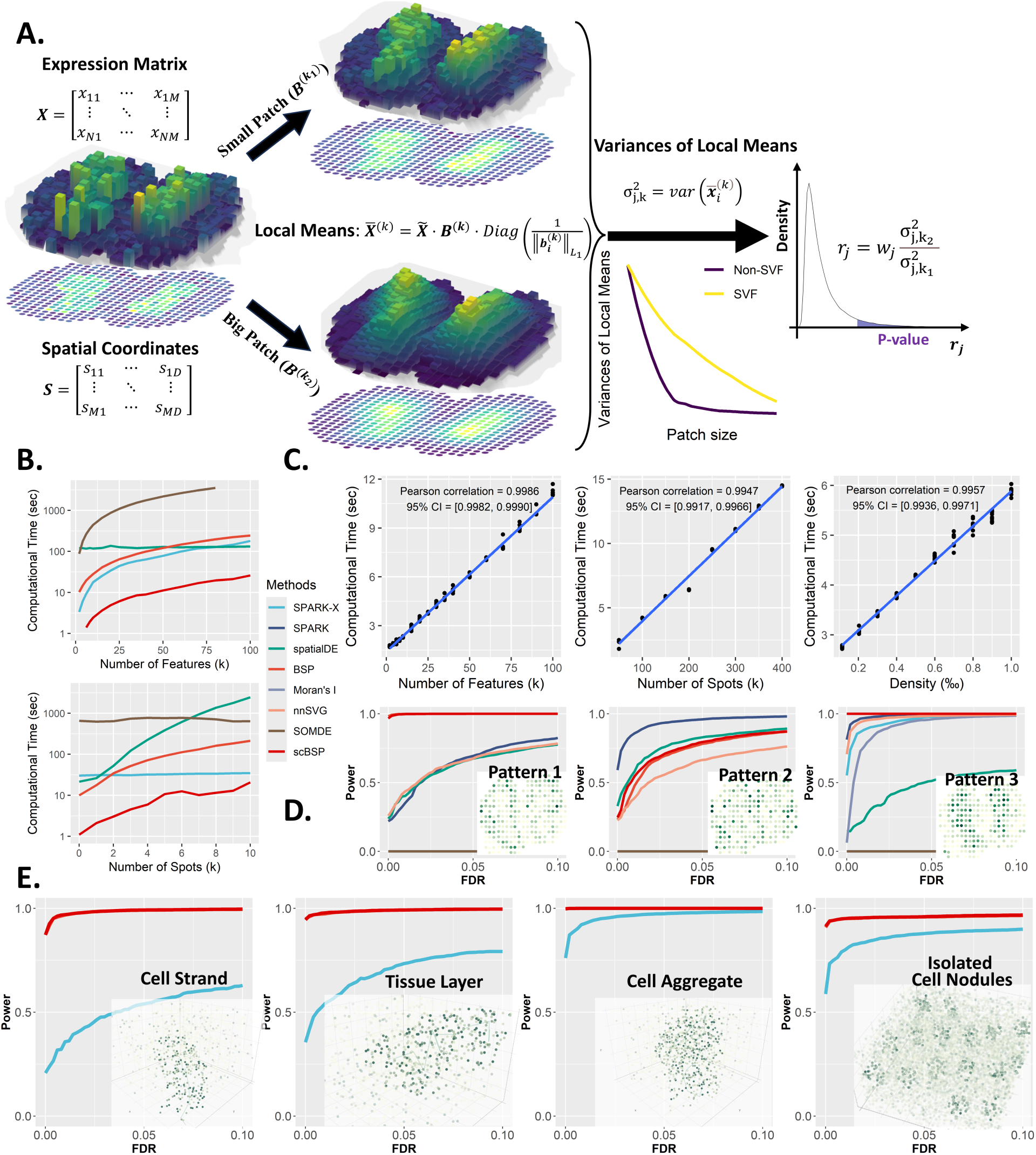
scBSP’s method and performance. **A:** Schematic of scBSP. The input comprises an expression matrix together with spatial coordinates. **B:** Computational time (y-axis) for analyzing spatial omics data comprising 20,000 features across 3,000 spots, varying number of features (top) and number of spots (bottom) while keeping the other constant. **C:** Computational time (y-axis) of scBSP on the high-resolution spatial omics data (run n = 10 times on a single processor core) with a varied number of features, spot count, and data density (x-axis). **D**: Comparisons of statistical power (y-axis) against false discovery rate (x-axis) on 2D simulations with spatial patterns I-III in mouse olfactory bulb (left to right) as defined in SpatialDE and SPARK. **E:** Comparisons of statistical power (y-axis) against false discovery rate (x-axis) on 3D simulations with Curved Cell Strand, Tissue Layer, Irregular Cell Aggregate, and Isolated Cell Nodules patterns (left to right).

For the spatial omics data at the spot level, scBSP outperformed all other tools when analyzing the data with 2,000 to 100,000 features on 3,000 spots (**Figure 1B**). Moreover, scBSP’s running time linearly increased with the number of features, highlighting the application on whole transcriptomic profiling and spatial omics data. For the samples with 20,000 features on 500 to 10,000 spots, scBSP also demonstrated lower computational time than other tools, particularly with an increase in spot count. While it consumes more memory than SPARK-X, scBSP generally requires less than 2GB of memory usage, making it suitable for processing spatial omics data with tens of thousands of features from thousands of spots on personal laptops (**Supplementary Figure 1**). Notably, nnSVG, Moran’s I and SPARK failed to process any data within the maximum allowed time (**Supplementary Table 1**),

We further explored computational efficiency on high-resolution spatial data comprising 20,000 features on 100,000 spots with a data density of 0.0005, and a serial of simulated data varying feature number, spot number, and density (as detailed in the Methods section). scBSP and SPARK-X were the only tools capable of processing such a large sample within hours on a desktop computer. Notably, scBSP analyzed most samples within 10 seconds, outpacing SPARK-X (**Supplementary Figure 2**). Similar to simulations with relatively smaller samples, scBSP’s running time was more sensitive to spot count and data sparsity but less to feature count, highlighting the computational efficiency of scBSP when dealing with high-resolution data from whole transcriptome sequencing methods or multi-omics data characterized by high expression sparsity. The memory usage of scBSP was higher than SPARK-X but still less than 2GB (**Supplementary Figure 1**). scBSP was also tested on mainstream high-resolution SRT technologies of 10x Visium, Stereo-seq, HDST, 10x Xenium, and CosMX. **Supplementary Table 2** details its running time and memory consumption.

scBSP’s computational efficiency lies in its linear scalability with the number of features, spots, and data density, which is crucial for detecting SVFs in high-resolution spatial omics data. We recorded computational time and peak memory usage of scBSP on high-resolution simulations with 10 replicates. While peak memory usage increased gradually at lower feature counts and data densities due to distance calculation requirements (with 100,000 spots in the high-resolution simulations), both runtime and peak memory usage scaled linearly with increasing feature count, spot count, and data density (**Figure 1C**, **Supplementary Figure 1**).

To ensure comparable effectiveness to the original BSP algorithm, we applied scBSP to simulations adopted from previous studies^32,34,35^, comparing performance with BSP and other SVG detection methods. Statistical power was assessed under a wide range of false discovery rates (FDR) to fairly evaluate model performance, which considered the discrepancies in the distribution of calibrated p-values across each method. In 2D simulations based on mouse olfactory bulb data with 260 spots, scBSP exhibited comparable statistical power to BSP across all three spatial patterns (**Figure 1D**). Notably, while SOMDE failed to detect all three spatial patterns, SPARK-X and Moran’s I also missed the first two patterns, leading to poor statistical power (**Figure 1D**). We also evaluated scBSP’s performance under varied signal strengths (**Supplementary Figure 3**) and noise levels (**Supplementary Figure 4**). We observed consistent performance of detection capacity in most scenarios.

In 3D simulations with a larger number of spots ranging from 2,000 for continuous spatial patterns to 9.000 for isolated patterns, scBSP demonstrated equivalent performance to BSP for all continuous and discrete spatial patterns (**Figure 1E**), surpassing SPARK-X. We varied pattern sizes, signal strengths, and noise levels (**Supplementary Figure 5-8**), and continuously observed robust detection capabilities across scenarios. Additionally, we assessed model performance with varied dropout rates (**Supplementary Figure 9**) and 3D data featuring inconsistent inter-plane and within-plane spatial resolution (**Supplementary Figure 10**). From these simulations, scBSP consistently exhibited statistical power comparable to BSP, outperforming competitive methods in all scenarios.

### scBSP effectively and efficiently identifies spatially variable features in spatial multi-omics data

In addition to analyzing SVGs in spatial transcriptomics data, we demonstrated scBSP’s capabilities in effectively and efficiently identifying SVFs in spatial ATAC-seq datasets. We evaluated the computational efficiency on two mouse spatial omics datasets to identify the most spatially variable ATAC peaks and compared the results with competitive methods of SPARK-X and spaLDVAE. Using the same quality protocols adopted by former studies ^36^, the first dataset is a mouse embryonic MISAR-seq dataset^37^ including fewer than 2,000 spots and approximately 47,000 ATAC peaks, while the second is the P22 mouse brain dataset^38^ containing over 9,000 spots and around 121,000 peaks (**Figure 2A**). All methods were tested on a Linux Ubuntu workstation with a 5.6 GHz Intel Core i9-13900 CPU and 32 GB RAM.

**Figure 2:**
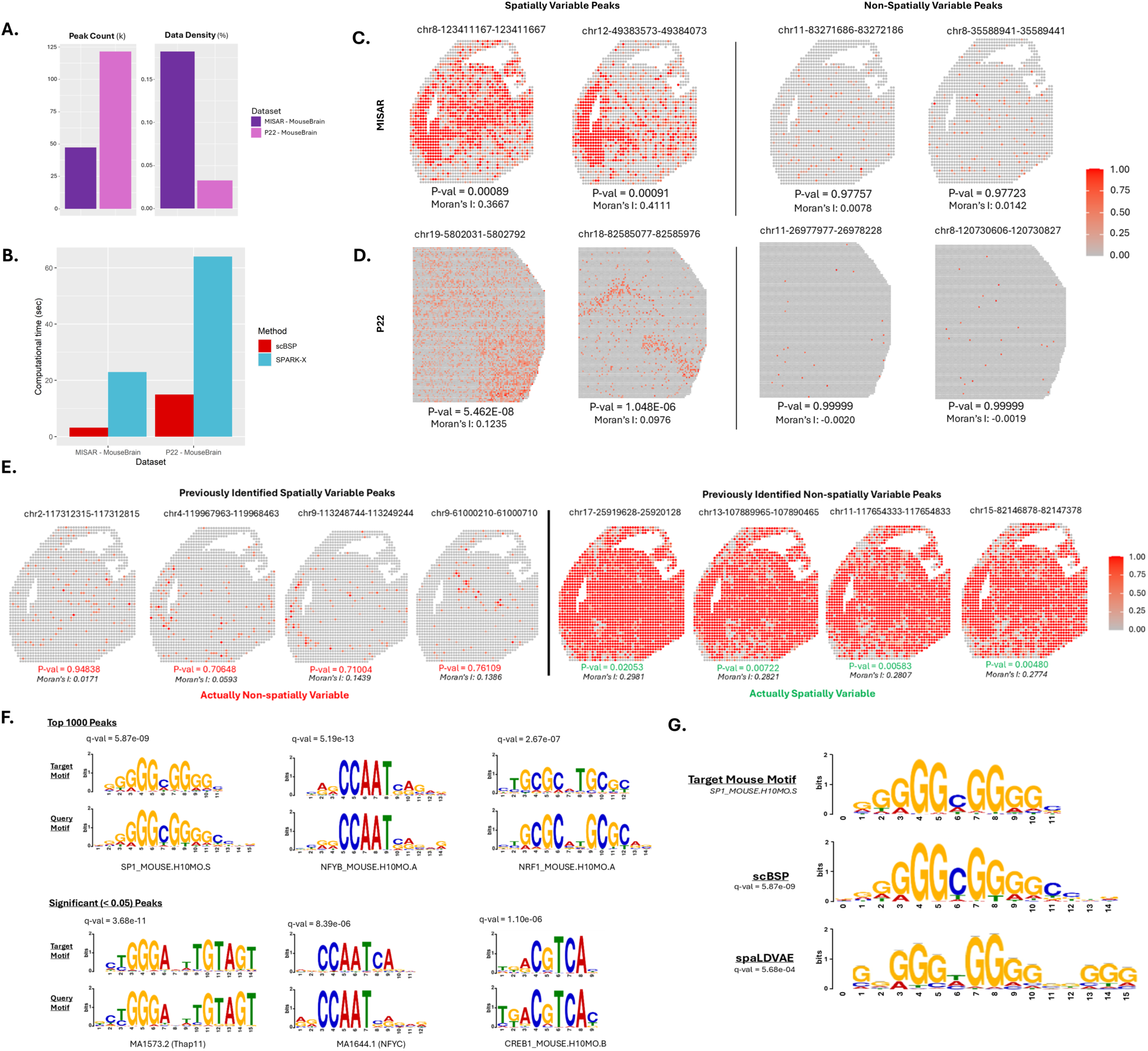
Analysis of spatial omics on mouse brain data. **A:** Peak count and data density of spatial ATAC-seq dataset. **B:** Computational time of scBSP and SPARK-X on each dataset. **C:** The ATAC counts of the top two significant and insignificant spatially variable peaks from scBSP on the MISAR mouse brain dataset. **D:** The ATAC counts of the top two significant and insignificant spatially variable peaks from scBSP on the P22 mouse brain dataset. **E:** Spatially variable ATAC-seq peaks highlighted in previous studies, which were not identified by scBSP (left), and ATAC-seq peaks identified by scBSP, which were not identified in previous work (right). **F:** MEME-ChIP identified sequence motifs from the top 1,000 spatially variable peaks (top panel) and all significant peak regions (bottom panel) detected by scBSP. For each alignment, the upper section displays known motifs in the mouse genome, while the lower section shows the query motifs identified from the significant peak regions. The q-values for a match between the known and query motifs were provided. **G:** Alignment comparison of SP1_Mouse between scBSP and SpaLDVAE, along with corresponding q-values.

We observed that scBSP outperformed competing tools in computational time on both datasets while maintaining reasonable memory usage (**Figure 2B**, **Supplementary Table 2**). Notably, spaLDVAE failed to complete either dataset within 24 hours. Biologically, scBSP demonstrated biological relevance in identifying spatially variable peaks. Using a unified significance threshold of 0.05, scBSP identified 8,337 out of 47,000 in the MISAR-seq dataset and 17,739 out of 121,000 spatially variable peaks in the P22 dataset, respectively. In contrast, SPARK-X identified 43,000 peaks as statistically significant, which is over 91% of the total peaks in the MISAR-seq dataset, potentially overestimating the results. Following the protocol used by spaLDVAE, we visualized the top four most and least spatially variable peaks identified by scBSP in **Figure 2C-D** and **Supplementary Figure 11**. These identified spatially variable peaks were also supported by positive values from the classical spatial autocorrelations measured using Moran’s I, while non-spatially variable peaks showed

isolated, random regions of high accessibility with Moran’s I value near zero (**Figure 2E**). We found the opposite trend between spatially and non-spatially variable peaks compared to results from SpaLDVAE (**Figure 2E**), which contradicts the definitions of Moran’s I on spatial autocorrelation.

Using MEME-ChIP, we further analyzed significant peak regions and identified enriched regulatory motifs among the spatially variable peaks ^39^. For both scATAC-seq mouse datasets, we evaluated scBSP results using only the top 1,000 spatially variable peaks and all significant peaks with p-values less than 0.05. Regulatory motifs identified from both scenarios demonstrated strong alignment and enrichment for binding sequences associated with specific, well-characterized mouse transcription factors (**Figure 2F**). For both the MISAR and P22 datasets, one of the identified motifs, ‘NFYB_MOUSE’ (nuclear transcription factor-Y beta), was strongly correlated with other ‘NFY’ variants known to bind the CCAAT element (q-value of 5.19e-13 and 8.39e-6 for each scenario, respectively)^40^. Another identified motif, ‘NRF1_MOUSE’ (nuclear respiratory factor 1), is particularly significant in brain regions, as its conditional deletion has been shown to cause severe neurodegeneration^41^. Notably, scBSP’s results demonstrated better alignment with the known motif ‘SP1_MOUSE’ than spaLDVAE (**Figure 2G**).

### scBSP robustly detects SVGs among different SRT technologies

Different SRT technologies reveal the inner characteristics of the tissue samples from different perspectives, varying resolution, sensitivity, specificity, and data sparsity. We evaluated the performance of existing methods in different SRT technologies. SVGs identified by scBSP are conservative among these different SRT technologies, given the variance in expression in the spatial spaces.

#### Analysis of high-resolution SRT data from mouse olfactory bulb

To illustrate the practical use of scBSP on high-resolution SRT data, we processed three high-resolution mouse olfactory bulb data collected from different sequencing techniques (Sections 1 and 2 from Stereo-seq and one from HDST). On a desktop equipped with Intel Core i9-13900 and 32GB memory, scBSP took 53.52 seconds and 65.37 seconds to process the Stereo-seq data with 26,145 genes measured on 107,416 spots and 23,815 genes on 104,931 spots, while SPARK-X took 51.14 seconds and 46.47 seconds, respectively. For the HDST data with more spots and increased data sparsity, scBSP only took 5.95 seconds to process the whole data, which consists of 19,950 genes measured on 181,367 spots, while SPARK-X took 33.16 seconds (**Supplementary Table 2**).

The detected SVGs from scBSP are consistent across the three olfactory bulb datasets from different sequencing technologies of Stero-seq and HDST. Specifically, scBSP identified 2,139 SVGs out of 26,145 genes in the first section from Stereo-seq, 2,057 SVGs out of 23,815 genes in the second section, and 2,292 SVGs out of 19,950 genes in HDST data (p-value<0.05). Overall, scBSP detected a total of 3,080 SVGs, with 1,363 (44.25%) of them shared among all three datasets (**Figure 3A**). In contrast, only 1.23% were shared over the total number of SVGs identified by SPARK-X. We further assessed the similarity between the identified SVGs from each pair of data using the Jaccard Index (JI). Specifically, for scBSP, the similarity score between two sections of Stereo-seq data was notably high at 0.75, declining to 0.54 and 0.48 when comparing Stereo-seq data with HDST data. In contrast, the similarity score for SPARK-X was 0.59 between two sections of Stereo-seq data but decreased rapidly to 0.02 and 0.01 when comparing Stereo-seq data with HDST data.

**Figure 3:**
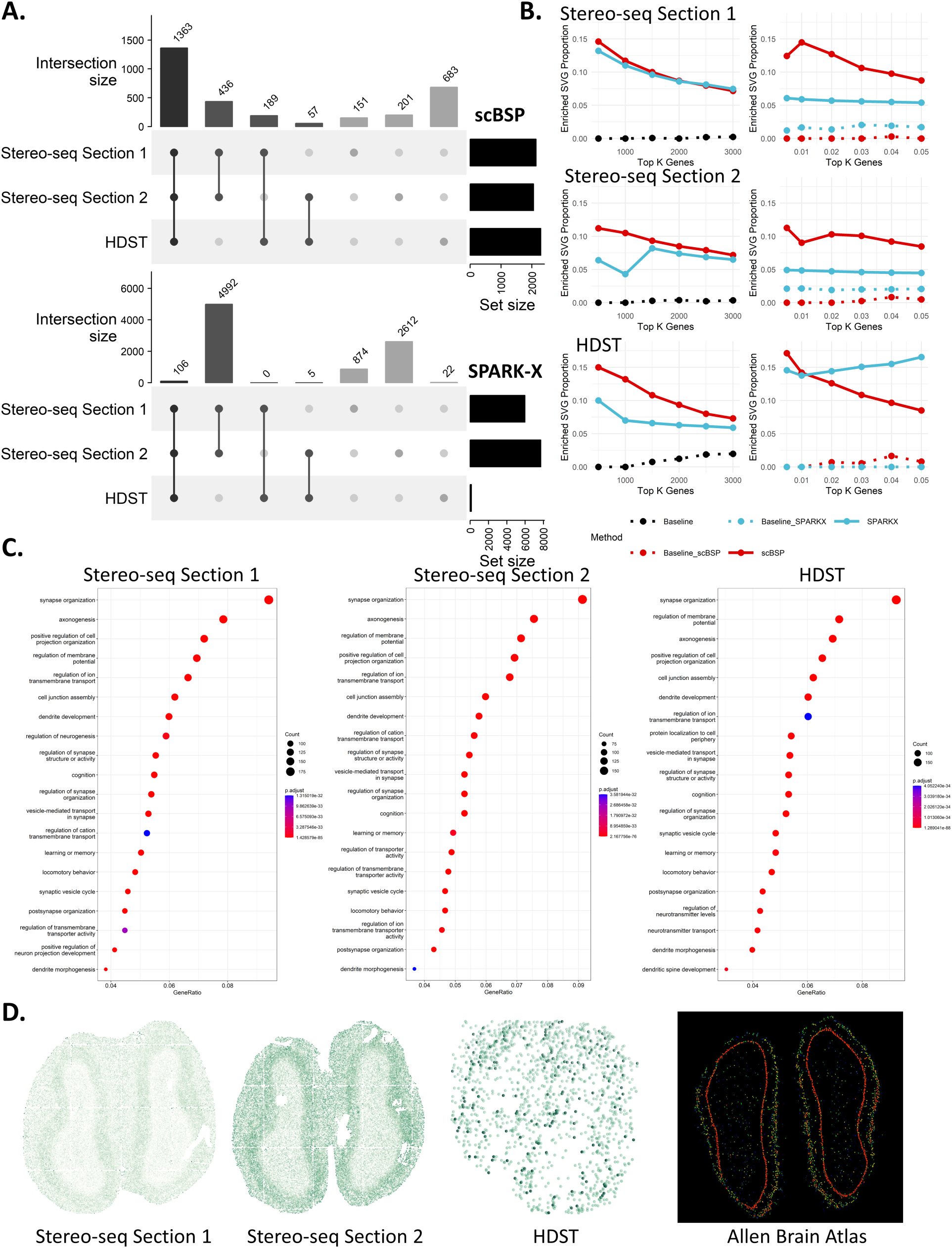
Analysis of mouse olfactory bulb data. **A:** Upset plot of identified SVGs from scBSP (upper) and SPARK-X (bottom) on three mouse olfactory bulb data. **B:** Proportions of enriched SVGs over top K genes (left) and proportions of enriched SVGs over the identified SVGs under varied type I error thresholds (right) on each data. **C:** Enriched gene ontology terms are based on scBSP-identified SVGs in each piece of data. **D:** Expression pattern of gene Ptprd in each dataset and Allen Brain Atlas.

We further investigated the proportions of enriched SVGs relative to the total identified SVGs across varying Type I error thresholds, and their proportions relative to the top K genes (ranging from 500 to 3,000) ranked by p-values (**Figure 3B**). Enriched SVGs were delineated based on enriched GO terms derived from GO enrichment analysis, utilizing either the top K genes with the lowest p-values from scBSP or SPARK-X (left side in **Figure 3B**), or using the genes with p-values below a specified threshold (right side in **Figure 3B**). While proportions over the top K genes were similar between scBSP and SPARK-X on Stereo-seq data, proportions based on the Type I error thresholds ranged from 8% to 15% for scBSP, surpassing SPARK-X (4% to 6%). However, proportions over the top K genes from SPARK-X were slightly lower than scBSP in HDST data, which may have resulted from SPARK-X only detecting 133 SVGs at a 0.05 threshold. This underscores the accuracy and robustness of scBSP in practical applications, particularly when using p-value thresholds.

The Gene Ontology (GO) enrichment analysis results exhibited high consistency on scBSP- identified SVGs across the three olfactory bulb datasets. The top seven enriched GO terms (synapse organization, axonogenesis, positive regulation of cell projection organization, regulation of membrane potential, regulation of ion transmembrane transport, cell junction assembly, and dendrite development) remained consistent across all three datasets from Stero-seq and HDST (**Figure 3C**). The olfactory bulb is the primary center in the processing of olfactory information, as it receives, filters, and transmits olfactory signals from the sensory neurons of the olfactory epithelium to one or more cortical olfactory centers, which highly corresponds to the organization of synapses shown in **Figure 3C** ^42,43^. In addition, the olfactory bulb is one of the few neurogenic regions that continues to be active throughout life. This is associated with axonogenesis, dendritic development, various aspects of learning, and memory performance as illustrated in **Figure 3C** ^44^. A representative gene uniquely identified by scBSP, Ptprd (**Figure 3D**), remained distinguishable even in HDST data characterized by a higher dropout rate (p-values of 6.26e-4, 7.71e-4, and 1.73e-05 for Stereo-seq data section 1, Stereo-seq data section 2, and HDST data, respectively). Ptprd is known to be intricately involved in axon guidance, synapse formation, and cell adhesion within the mouse brain, which has been found to regulate NMDAR-mediated postsynaptic responses in neural circuits that are spatially linked to asynchronous release sites in recent studies ^43,45,46^. Overall, SVGs identified by scBSP accurately characterized (highly associated with) the biological functions in the mouse olfactory bulb tissue data.

#### Analysis of SRT data from wholhe mouse brain

We further applied scBSP to analyze five whole mouse brain samples with sequencing technologies at a relatively lower resolution. Four sagittal datasets were collected using the 10x Genomics Visium platform, while the coronal dataset was obtained from Stereo-seq sequencing. On the desktop, scBSP demonstrated computational efficiency by processing each 10x Visium dataset in an average of 5 seconds, and 25.11 seconds for the Stereo-seq data. In comparison, SPARK-X required 36 seconds per 10x Visium dataset and 47.19 seconds for the Stereo-seq dataset (**Figure 4A**). In contrast, scBSP took longer than SPARK-X when processing the Xenium and CosMx SRT data where genes are pre-selected (**Figure 4A**).

**Figure 4:**
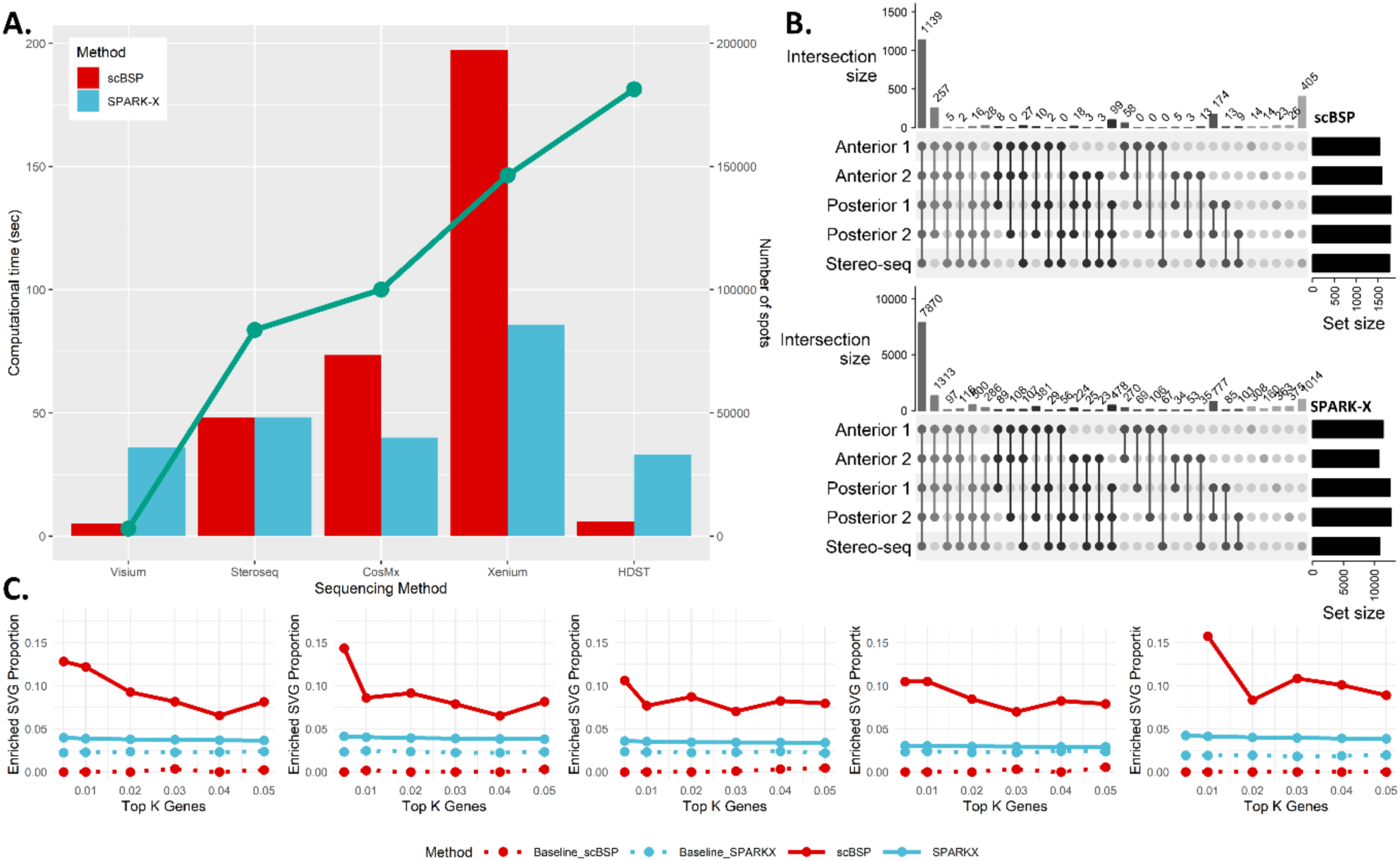
Analysis of mouse brain data. **A:** Computational time of scBSP and SPARK-X on real data. The bar charts illustrate the running time (left y-axis) on each data (x-axis). The green line denotes the number of spots in each dataset (right y-axis). **B:** Upset plot of identified SVGs from scBSP (upper) and SPARK-X (bottom) on five mouse brain data. **C:** Proportions of enriched SVGs over top K genes (left) and proportions of enriched SVGs over the identified SVGs under varied type I error thresholds (right) on each data.

Similar to observations in olfactory bulb data, the identified SVGs from scBSP were consistent across various datasets, particularly exhibiting high consistency between the two anterior sections (JI=0.92) and two posterior sections (JI=0.94) (**Figure 4B**). Moreover, the enriched pathways detected by scBSP were very close across datasets, which were related to the biological processes in the mouse brain (**Supplementary Figures 12-16**). We also examined the proportions of enriched SVGs over the identified SVGs across varied type I error thresholds (**Figure 4C**). The threshold-based proportion of scBSP was consistently higher than SPARK-X, highlighting the application of scBSP as the SVGs were typically selected based on a given threshold of type I error rate in practice. In summary, scBSP efficiently and robustly identified biologically meaningful SVGs in SRT invariant to the sequencing technologies and resolutions.

#### Analysis of single-cell 10x Xenium SRT data from human kidney

We analyzed a kidney dataset generated using the 10x Xenium platform, which included 12 kidney samples: 4 from healthy controls, 4 from patients with lupus nephritis, and 4 from patients with chronic kidney disease (CKD) ^47^. Each sample contained over 25,000 cells and 300 genes (**Supplementary Table 3**). Initially, we applied scBSP to identify SVGs among the 300 genes within each sample. Subsequently, we conducted a meta-analysis using Stouffer’s method to calculate combined p-values across the 4 samples within each condition (**Supplementary Table 4**), selecting genes with meta-p values below 0.05. These significant SVGs were further analyzed using hierarchical clustering to group their expression counts (**Supplementary Table 5**).

In each condition, we identified a gene cluster significantly smaller than others, comprising only 3-5 genes, in contrast to larger clusters containing over 30 genes (**Figure 5A**). We then identified overlaps in SVGs across the different conditions (**Figure 5B**). We observed that VIM (vimentin) and AQP1 (aquaporin-1) were consistently significant across all three conditions, with meta-p values of 9.03e-141, 0, and 1.14e-121 for VIM, and 0, 1.79e-193, and 8.43e-14 for AQP1 in lupus nephritis, CKD, and control samples, respectively. Analyzing the full SVG sets showed that most genes were found in all three condition SVGs, but post-clustering analysis revealed that VIM was exclusively found in the small cluster associated with CKD and lupus nephritis, whereas AQP1 was specific to the control group’s cluster, indicating distinct spatial expression patterns between diseased and healthy conditions (**Figures 5C-5D**). VIM is a known marker of epithelial to mesenchymal transition observed in kidney disease ^48^. AQP1, or aquaporin-1, is a water transport channel and a canonical marker found in healthy proximal tubule cells. A similar pattern was also observed for DDX5, B2M, and TMSB4X in CKD/lupus nephritis, and for GATM in the control group (**Supplementary Figure 17**).

**Figure 5:**
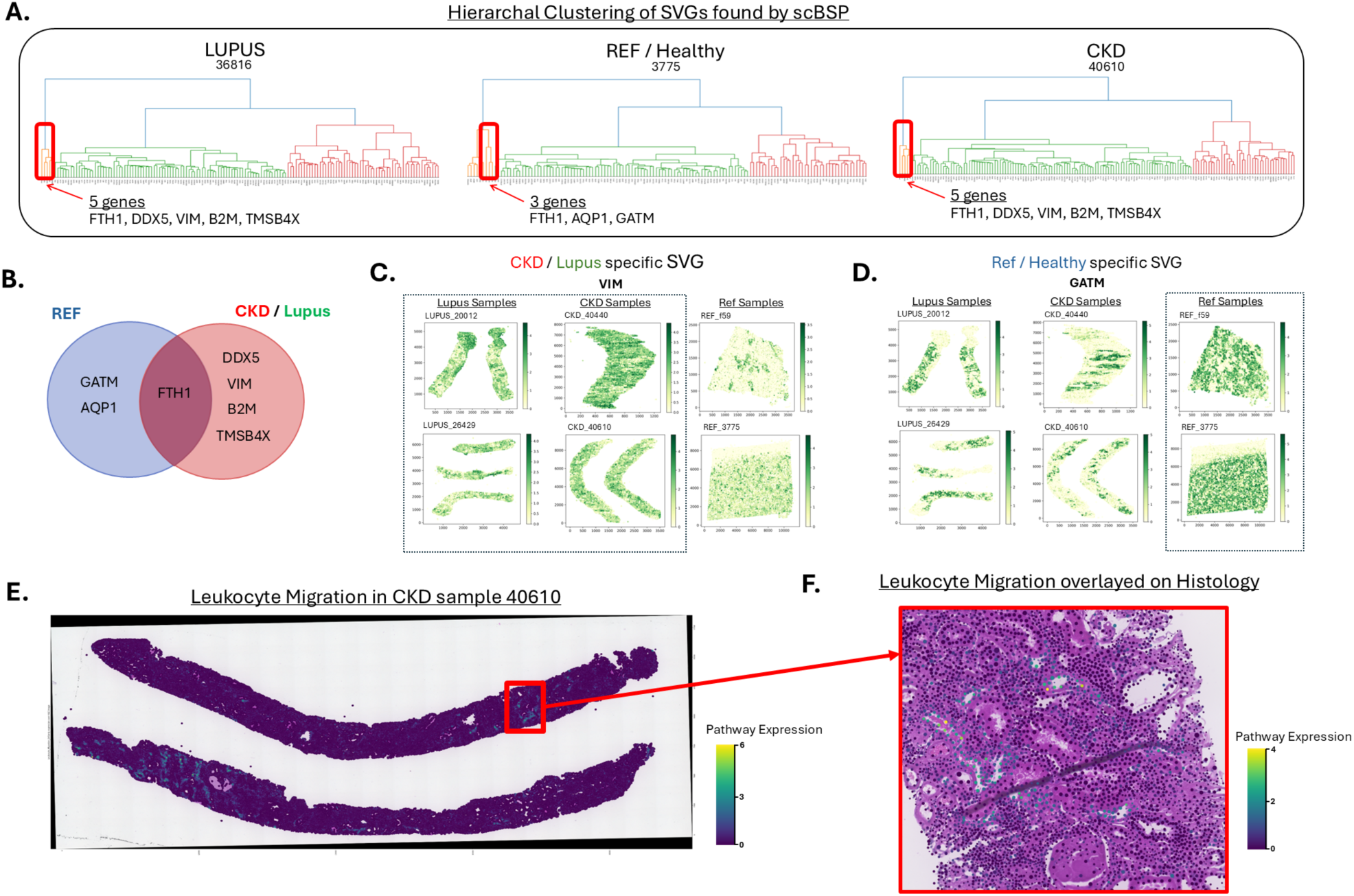
SVG analysis on 10x Xenium Kidney data using scBSP. **A:** Hierarchical clustering analysis on SVGs. The highlighted cluster displays the unique cluster in each condition. **B:** Venn diagram of the unique cluster from each condition, showcasing the overlap between the SVGs in Reference and CKD/Lupus. **C:** Gene expression of the 10x Xenium kidney samples. This group plots the gene expression of VIM, a CKD/Lupus-specific SVG. **D:** This group plots the gene expression of AQP1, a reference-specific SVG. The difference in expression is observed in the CKD/Lupus vs. reference samples. **E:** Sample 40610 – CKD, H&E image overlapped with pathway expression of leukocyte migration, showing the whole sample’s pathway expression on both the full samples. **F:** Cropped section of sample 40610, which is overlayed on top of the H&E image. This alignment allows cross referencing of histology with upregulated immune expression from the pathway.

For larger clusters in each condition, unique SVGs in each condition were analyzed by Gene Ontology Enrichment analysis and Pathway Enrichment analysis using clusterProfiler (**Supplementary Figures 18-19**). In Lupus, out of 93 genes in the largest cluster, 43 SVGs were unique to Lupus compared to reference SVGs. Compared with all 93 SVGs contributing 16/93 genes or 17.2% to the ERK1 and ERK2 pathway, unique SVGs contributed 10/43 genes or 23.3% of the gene set to this crucial Lupus pathway. Similarly, the largest reference cluster of 78 genes found ERK1 and ERK2 cascade as a significant pathway (q-value 0.0045), but when focusing on the 28 unique SVGs for the condition, this pathway is no longer found as significant (q value 0.3040). Leukocyte migration is also an important pathway that plays a role in CKD’s progression ^49^. This pathway is significantly enriched in CKD’s 58 unique SVGs set (q value 7.37e-03) but is not identified as significant within the reference group’s 20 unique SVGs. These results suggest unique SVGs within different conditions may amplify and reveal some biological mechanisms between different diseases.

To further uncover the differences in pathway expression between the reference sample F59 and CKD sample 40610, the pathway expression of Leukocyte migration was quantified by subtracting the average expression of the pathway gene set with aggregated expression of a randomly sampled set of control genes using Scanpy’s *score_gene* function ^50^. Incorporating the pathway expression in the reference sample, Leukocyte migration was localized in specific areas within the spatial context upon aligning with H&E histology images (**Supplementary Figure 20**). The expression of the Leukocyte migration pathway was more widespread across the whole CKD sample than the reference sample. Cells that highly expressed this pathway were surrounded by cells with moderate to low expression (**Figure 5E**).

## DISCUSSION

The dawn of spatial omics enables new challenges and opportunities to explore the relationships between the structure and functionalities. scBSP is developed as a computational efficiency package available in R and Python to identify SVFs in high-resolution spatial omics data. The advent of high-resolution spatial sequencing techniques has seen a drastic increase in the number of spots per sample, escalating from hundreds to millions over the past decade. This surge in resolution poses a challenge for most of the existing tools, many of which require hours to days to process a single SRT sample. scBSP addresses the critical need for computational tools capable of detecting SVFs in high-resolution spatial multi-omics data, particularly those characterized by high sparsity. In our simulation and case studies, scBSP demonstrates remarkable efficiency by processing samples with hundreds of thousands of spots and features in seconds to minutes on a typical desktop. For instance, on a typical desktop, scBSP processed the high-resolution SRT data from HDST sequencing comprising 181,367 spots in 5 seconds, which is the fastest tool available to our knowledge for handling such high-resolution, sparse data while maintaining high accuracy. Notably, the scBSP Python library is the only tool for handling such high-resolution data within reasonable computational resources. This improved computational efficiency accelerates SVF inference in Python and makes spatial transcriptomics accessible to more researchers, driving faster insights into spatial expression and accessibility patterns across various biological processes and allowing for the implementation of well-developed deep learning frameworks among the Python community.

scBSP implements sparse matrix operation to take advantage of algorithm optimization and compiler optimization in contemporary computer architectures. The scBSP R package is conveniently hosted on the Comprehensive R Archive Network (CRAN), requiring no external software dependencies other than R packages, ensuring straightforward installation and usability across all major operating systems. Similarly, the scBSP Python library is readily downloadable and installable via the Python Package Index (PyPI), offering both a full version reliant on Microsoft Visual C++ for approximate nearest neighbor search and a lightweight version with no dependencies beyond Python libraries. Additionally, the scBSP R package boasts a high level of interoperability with other tools, accepting inputs in the form of Seurat objects, thus seamlessly integrating with a wide array of analysis and visualization functions within the Bioconductor ecosystem.

Furthermore, scBSP’s non-parametric, spatial granularity-based model offers a fast, accurate, and robust solution for SVG detection in high-resolution SRT data. Simulations demonstrate scBSP’s superior performance in most scenarios, exhibiting robustness to varying signal strengths, noise levels, and dropout rates. Moreover, when applied to high-resolution olfactory bulb datasets collected from different sequencing techniques, the identified SVGs from Stereo-seq and HDST data exhibit high consistency (JI = 0.54) despite a significant increase in the data sparsity, while the number of identified SVGs from SPARK-X decreased from 5972 to 133 (JI = 0.02). This is potentially because the spatial granularity-based model does not make assumptions about the underlying distributions of expressions, which may not hold in the complex biological environment and the expression data with high dropout rates. This comparison across SRT samples highlights scBSP’s reliability and robustness, which are essential for obtaining meaningful insights across experiments and sequencing technologies. Notably, this characteristic of scBSP will benefit more with the advancement of new high-throughput, high-sensitivity sequencing technologies in the future. Additionally, scBSP’s ability to detect enriched SVGs and their consistent enrichment in gene ontology terms across datasets further validates its effectiveness in capturing biologically relevant information, which could be vital in assisting pathologists in identifying the pathogenesis of different diseases.

scBSP also demonstrates its robustness on spatial ATAC-seq data, not compromising computational efficiency when running over a hundred thousand chromatin peaks on a desktop. Downstream analysis of spatially variable chromatin regions showed that scBSP can identify peaks with high spatial autocorrelation, giving less importance to those with more random spatial distributions. Motif analyses also reveal that top spatially variable peaks in both mouse samples found regions vital to the brain region, amplifying the method’s biological findings as well. Additionally, the application of scBSP on high-resolution 10x Xenium SRT data demonstrates its ability to identify key SVGs in different conditions. Further analysis of condition-unique SVGs found by scBSP shows they successfully highlight significant SVPs within the conditions. With a lack of robust SVP analysis methods, this proof of concept by mapping key pathways found using scBSP onto sample histology images could be a key tool in aiding pathologists in discovering novel SVGs and spatially variable pathways that contribute to the pathogenesis of kidney disease.

While the scBSP package offers significant advancements in SVF detection on high-resolution spatial omics data, it still presents some limitations. The Python version depends on external software (Microsoft Visual C++ 14 or higher) for nearest neighbor detection, with a Ball Tree alternative offered to ensure compatibility. However, this alternative may slightly reduce efficiency and introduce minor methodological differences. Additionally, scBSP’s computational time and memory usage are sensitive to data sparsity. While it outperforms SPARK-X on the benchmarks, it may require more time and memory than SPARK-X for high-resolution SRT data with higher data density and lower number of features, such as Xenium and CosMx SRT data where genes are pre-selected. In the future, we will delve into scBSP’s applications across a range of high-resolution spatial omics case studies, including analyses of the tumor microenvironment, Alzheimer’s disease, and kidney research. We also continue to explore scBSP’s potential with other spatial omics data at the cellular or subcellular level, such as metaFISH, spatial CITE-seq, and spatial CUT&Run-seq, and apply it to high-resolution spatial-temporal studies such as embryogenesis in mouse embryos using Stereo-seq platforms.

## METHODS

### Overview of scBSP workflow

scBSP is an open-source R package and a corresponding Python library that implements a spatial granularity-based algorithm for identifying SVFs in dimension-agnostic and technology-agnostic spatially resolved transcriptomics and multi-omics data. A pair of patches is defined for each spot in the spatial data, comprising neighboring spots within a small and large radius. The average expression within each patch is calculated, reflecting the local expression levels at various spatial granularities. The variance of local expressions is computed across all spots for each given radius. The key concept of this approach is to capture the rate of change in variance of local means as the granularity level increases, measured by the variance ratio with different patch sizes (**Figure 1**).

Given the substantial number of spatial locations and the high sparsity of the expression matrix in high-resolution SRT, scBSP adapted the BSP algorithm for expression rescaling, patch determination, and local expression calculation (**Supplementary Figure 21**) with lower time complexity and space complexity (**Table 1**). Specifically, for an SRT sample with 𝑀 spots and 𝑁 features, the 𝐷-dimensional coordinates of spot 𝑖 are denoted as a 𝐷-vector 𝒔*_i_*, where 𝑖 = 1, …, 𝑀, and the corresponding coordinates matrix of all spots are denoted as a 𝑀 × 𝐷 matrix 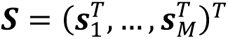. For a spot 𝑖_0_ in the sample, we define the patch as the set of neighboring spots within the radius 𝑅*_k_*, denoted as a binary M-vector 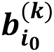, where 𝑘 = 1,2 for small and big radius, respectively, 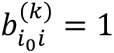 if 𝑑𝑖𝑠𝑡(𝑖_0_, 𝑖) < 𝑅*_k_* and 𝑖 ≠ 𝑖_0_. To avoid an empty patch when a spot has no neighboring spots within the given radius, we assign the spot itself to the patch to represent the local expression levels. That is, 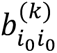 is assigned the value of 1 if and only if 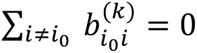. The patches of all spots in the SRT sample are thereby denoted an 𝑀 × 𝑀 binary patch matrix, 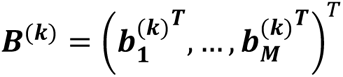, for each given radius 𝑅*_k_* . We consider paired big-small patches with radii 𝑅_1_ and 𝑅_2_, where 𝑅_1_< 𝑅_2_, with the default values as one and three units as demonstrated in BSP ^32^.

**Table 1:** Time and space complexity of BSP and scBSP. M: number of spots; N: number of features; S: data sparsity (proportion of non-zero values in the expression matrix). The Approximate Nearest Neighbour Search in Python is based on HNSW implementation. Note: practical performance may vary from theoretical analysis due to factors such as compiler optimizations and specific package implementations.

| Process | BSP |  |  | scBSP |  |  |
| --- | --- | --- | --- | --- | --- | --- |
|  | Method | Time Complexity | Space Complexity | Method | Time Complexity | Space Complexity |
| **Overall** | - | $O(NM^2)$ | $O(NM)$ | - | $O(NM \log M)$ | $O(NM)$ |
| Data Pre-processing | Min-max Scaling | $O(NM)$ | $O(NM)$ | Maximum Absolute Scaling | $O(M)$ | $O(NM)$ |
| Patch Determination | Exhaustive Search | $O(NM^2)$ | $O(NM)$ | Approximate Nearest Neighbour Search* | $O(NM \log M)$ | $O(NM)$ |
| Expression Matrix Calculation | Dense Matrix | $O(M)$ | $O(NM)$ | Sparse Matrix | $O(SM)$ | $O(SNM)$ |
| Local Expression Calculation | Loop Operation | $O(NM)$ | $O(NM)$ | Matrix Operation | $O(SNM)$ | $O(SNM)$ |

To ensure an adequate number of spots captured by the pre-defined radii 𝑅_1_ and 𝑅_2_, the coordinates matrix is normalized based on the density of spots such that the average spot-to- spot distance to is slightly less than one unit. The rescaling function is defined as 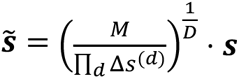, where 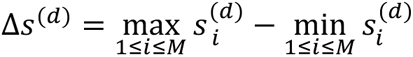 denotes the ranges of coordinates in each direction. The Euclidean distance between spots 𝑖_1_and 𝑖_2_, denoted as 𝑑𝑖𝑠𝑡(𝑖_1_, 𝑖_2_), is then calculated using approximate nearest neighbor search, which implements a K-Dimensional Tree with the threshold of 𝑅_𝑘_. For the SRT samples with cell counts ranging from thousands to hundreds of thousands, utilizing the approximate nearest neighbor search can achieve significantly faster running times with relatively small errors compared to the brute-force computation of all distances, as the time complexity is reduced from 𝑂(𝑀^2^) to 𝑂(𝑐 × 𝑙𝑜𝑔(𝑀)), where 𝑐 is a constant depending on the dimension and approximation error ^33,51^. The 𝑁 × 𝑀 raw expression matrix is denoted as 𝑿, where the raw expression level of feature 𝑗 in spot 𝑖 is denoted as 𝑥_𝑖j_, 1 ≤ 𝑖 ≤ 𝑀, 1 ≤ 𝑗 ≤ 𝑁. Considering the sparsity of high-resolution SRT data, the expressions are rescaled to [0, 1] by features using the maximum absolute rescaling as

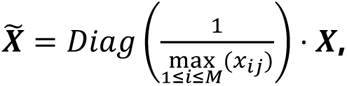

where 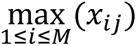 denotes the maximum expression level of feature 𝑗 and 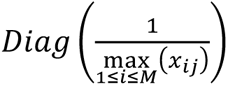 denotes the 𝑁 × 𝑁 diagonal matrix of the 𝑁-element vector 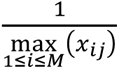. The matrix for averaged expression level 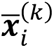 for a given radius 𝑅*_k_* at spot 𝑖, referred to as the local means, is calculated as

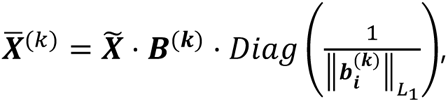

where 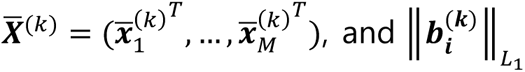, denotes the number of spots in the patch 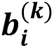. Subsequently, the variance of local means for feature 𝑗 is computed as 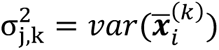 . We utilize the ratio of the variances of the paired local averaged expression levels between big and small patches, 𝑟_*i*_, to measure the velocity of changes in the variances of local means for feature 𝑗, defined as:

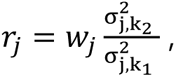

where 𝑤*_j_* is the weight to normalize the intrinsic expression variance within the feature with the maximum variance, i.e., 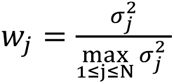 where 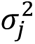 is the variance of raw expression levels of feature 𝑗 of all the spots in the sample, and 1 ≤ 𝑛 ≤ 𝑁. The distribution of 𝑟*_j_* is approximated with a lognormal distribution. The null hypothesis that a feature has no spatial pattern is thus reformulated as the ratio of a feature adhering to the fitted log-normal distribution. To tolerate the potential noise and long-tail deviations, a one-sided p-value is assigned to each feature if 𝑟*_j_* exceeds the upper tail of the fitted distribution at a probability of 100 × (1 − 𝛼)%, where 𝛼 refers to the significance level (usually set as 0.05).

We assume that a minority of features situated in the upper tail of the distribution exhibit spatial variability, while most features are non-SVFs. This proposition is particularly relevant in high-throughput SRT platforms like 10x Visium, which have an extensive feature repertoire exceeding thousands. Notably, in low-throughput SRT platforms where the gene count is limited (like MERFISH), a set of random features permuted across spatial locations is recommended to estimate the distribution of the test score for non-SVFs.

### Simulations

#### Simulation for benchmarks

To evaluate the accuracy of identifying SVFs across various scenarios, we conducted model performance assessments of scBSP and compared the statistical power with those obtained using the original BSP algorithm and other existing methods. This evaluation utilized simulated data from previous studies ^25,32,52,53^. The 2D simulations were based on mouse olfactory bulb data, comprising three spatial expression patterns observed across 260 spots (as detailed in the Results section). Each simulation included 1,000 simulated SVFs with recognized patterns from studies of SpatialDE and SPARK, alongside 9,000 non-SVFs generated via feature permutation without any discernible spatial expression pattern. Statistical power (true positive rates) was calculated using p-values derived from various methods, including basic spatial autocorrelation statistics (e.g., Moran’s I), SpatialDE, SPARK, SPARK-X, SOMDE, BSP, and scBSP, across a wide range of false discovery rates (FDR), accounting for the disparities in the distribution of p-values from each method. Ten replicates were included for each simulation to mitigate sample bias in the power analysis. Two key parameters were considered: signal strengths, quantified as the fold-changes (FCs) between average expressions in patterned and non-patterned regions (FC=3,4,5 for low, moderate, and high signal strengths, respectively), and noise level, defined by the dispersion parameters (𝜏_2_) in SPARK’s model (𝜏_2_=0.2,0.5,0.8 for low, moderate and high noise levels, respectively).

Similar to the 2D simulations, the 3D simulations comprised 1,000 simulated SVFs for each spatial pattern and 9,000 permuted features without any spatial patterns. Three continuous patterns (Curved Cell Strand, Tissue Layer, and Irregular Cell Aggregate) and one discrete pattern (Isolated Cell Nodules) were included, representing a strand-like arrangement of cell clusters (e.g., nerve strands), tissue layers, dense cell clusters (e.g., tumors), and discrete, spherical clusters (e.g., lymphoid follicles) (Figure 1E). The simulated data for continuous patterns consisted of 10 segments in the z-coordinate and 225 spots in each piece in the x- and y-coordinates, assuming the sample was cryo-sectioned with each section placed on an individual array without direct contact between array surfaces. The continuous patterns were designed using a set of spheres with center points generated through a random walk with a fixed step length of 2 units. Three patterns were constructed by controlling the number of directions of monotonic movements (Curved Cell Strand: monotonic in two directions; Tissue Layer: monotonic in one direction; Irregular Cell Aggregate: non-monotonic in any directions).

The simulations for discrete patterns were constructed on the data with 10 segments and 900 spots in each segment, where the x- and y- coordinates on each segment ranged from 0 to 30. The discrete spatial patterns were designed using solid spheres spaced 8 units apart. Sixteen center points were selected with a fixed z-coordinate of 5.5, while x- and y-coordinates were derived from a sequence of 3 to 27 at intervals of 8. Uniformly distributed noise ranging from −2 to 2 was added to each coordinate to introduce randomness.

The expression of SVFs was sampled based on whether the cell was inside or outside the pattern, distinguishing between marked and non-marked cells. For marked cells inside the pattern, we randomly selected feature expressions from the upper quantile of the feature expression distribution in the seqFISH data. For non-marked cells and those outside the pattern, we assigned feature expressions randomly from the seqFISH data. Non-SVFs were generated by permutating SVF’s expressions. Finally, random noise was added proportionally to all features’ average expression standard deviation. Three parameters were considered in the simulations, including pattern size, measured as the radius of the pattern (radius=1.5,2.0,2.5 for small, moderate, and large pattern size, respectively), signal strengths, quantified as the fold-changes between average expressions in patterned and non-patterned regions (FC=2.0,2.5,3.0 for low, moderate, and high signal strengths, respectively), and noise level, 𝜎, defined as the proportion to the averaged standard deviation of expressions in all features (𝜎 =0,1,2 for low, moderate, and high noise levels, respectively). Two additional simulations were incorporated to evaluate the model’s robustness in handling SRT data characterized by distinct inter-plane and within-plane spatial resolutions and the data with varied dropout rates, respectively. For this purpose, the simulated data were generated following the previously described procedure, with an additional step involving the multiplication of z-coordinates by 10 to simulate reduced inter-plane resolution. For the simulations with varied dropout rates, a random subset (10% to 30%) of spots was assigned a value of 0 to simulate the dropout events.

#### Simulations for computational efficiency

Recent studies showed that SPARK-X was the fastest method for SVG detection, while SOMDE is the second best in most cases but significantly slower than SPARK-X ^54,55^. However, the results were based on small samples with fewer than 40,000 spots. To fairly compare the model performance in the small and high-resolution SRT samples, respectively, we conducted comparisons of computational efficiency under two distinct scenarios: one involving a larger number of spots with higher data sparsity (referring to HDST data), and the other with a relatively smaller number of spots (referring to stxBrain data).

In the first scenario, two simulations were devised to assess computational efficiency across models. The first set varied the number of spots from 500 to 10,000 while keeping the number of features fixed at 20,000. The second set varied the number of features from 2,000 to 100,000 while maintaining a fixed number of spots at 3,000. Expression values were randomly sampled from the expression data of the first anterior of the "stxBrain" dataset in the "SeuratData" package. The metadata of the simulations were detailed in **Supplementary Table 1**.

In the second scenario, three simulations were conducted, each varying the number of spots, features, and data density, respectively. The first set involved a range of spots from 50,000 to 400,000, with a fixed number of features at 20,000 and a data density of 0.05. The second set varied the number of features from 2,000 to 100,000, with a fixed number of spots at 100,000 and a data density of 0.05. The final set varied the data density from 0.01% to 0.1%, with a fixed number of features at 100,000 measured across 100,000 spots. This approach was guided by the characteristics of the HDST data, which includes 19,950 genes measured across 181,367 spots, with a data density of 0.04%. Non-zero expression values were generated as follows: 𝐸𝑥𝑝∼⌊1 + 10 × 𝐵𝑒𝑡𝑎(1,5)⌋, where ⌊𝑥⌋ denotes the maximum integer that is not greater than x.

### Real Data Analysis

The 10x Genomics Visium mouse brain data were extracted from “stxBrain” data in the “SeuratData” package. For HDST data, we adopted the analysis protocol from SPARK-X. For studies involving Stereo-seq ^14^, 10x Xenium ^56^, and Cosmx datasets, we adhered to their respective original protocols. Specifically, for the 10x Xenium and Cosmx datasets, gene expressions were permuted to ensure a total gene count exceeding 10,000, facilitating sufficient null genes for scBSP and SPARK-X to determine the null distributions. The metadata of the simulations were detailed in **Supplementary Table 2**. Enrichment analysis was performed using the "clusterProfiler" package ^57^ in R. For spatial ATAC-seq analysis, cells and peaks were extracted directly from the h5 and h5ad files provided by the original studies. Both SPARK-X and scBSP were run on the full datasets. Results from scBSP were then subset to keep the statistically significant spatially variable peaks with p-values <0.05. These and the top 1000 peaks were then run through MEME-ChIP to find motifs for both MISAR and P22 datasets.

## DATA AVAILABILITY

All relevant data supporting the key findings of this study are available within the article and its Supplementary Information files. The stxbrain data can be downloaded with the “SeuratData” package in R. HDST data are available at Broad Institute’s single-cell repository with ID SCP420. The Stereo-seq data from the Mouse Organogenesis Spatiotemporal Transcriptomic Atlas (MOSTA) is available at https://db.cngb.org/stomics/mosta/download/. The 10x Xenium human breast cancer data and mouse brain data are available at https://www.10xgenomics.com/products/xenium-in-situ/preview-dataset, and https://www.10xgenomics.com/datasets/fresh-frozen-mouse-brain-replicates-1-standard.

The Cosmx dataset (Lung Cancer Slices) is available at https://nanostring.com/products/cosmx-spatial-molecular-imager/ffpe-dataset/nsclc-ffpe-dataset/. The simulation data are available in the Figshare database at https://doi.org/10.6084/m9.figshare.2418792360. The 10x Xenium Kidney data was provided by the Eadon laboratory, and will become publicly available soon. MISAR-seq dataset at https://figshare.com/articles/dataset/Spatial_genomics_datasets/21623148/5. P22 mouse brain dataset at https://zenodo.org/records/7879713.

## CODE AVAILABILITY

The Python library, “scBSP”, is available at https://pypi.org/project/scbsp/. A corresponding R package, “scBSP”, is available on R CRAN at https://cran.r-project.org/web/packages/scBSP/index.html. Project home page: https://github.com/CastleLi/scBSP/

Archived versions: https://zenodo.org/records/14768450 https://figshare.com/articles/software/scBSP_zip/26828332?file=48773866

## AUTHOR CONTRIBUTIONS

Conceptualization: JL, JW, QM, and DX; methodology: JL and JW; software coding: JL and YW; data collection and investigation: JL, MR, and JW; data analysis: JL, MR, YW, LS, and CX; pathology analysis: QG, YC, RF, and ME; software testing and tutorial: JL, and YW; manuscript writing, review, and editing: JL, MR, JW, QM, and DX.

## REFERENCES

1. Larsson, L., Frisén, J. & Lundeberg, J. Spatially resolved transcriptomics adds a new dimension to genomics. Nat Methods 18, 15–18 (2021).

2. Bressan, D., Battistoni, G. & Hannon, G. J. The dawn of spatial omics. Science 381, eabq4964 (2023).

3. Llorens-Bobadilla, E. et al. Chromatin accessibility profiling in tissue sections by spatial ATAC. Preprint at 10.1101/2022.07.27.500203 (2022).

4. Geier, B. et al. Spatial metabolomics of in situ host-microbe interactions at the micrometre scale. Nat Microbiol 5, 498–510 (2020).

5. Price, A. E. et al. A Map of Toll-like Receptor Expression in the Intestinal Epithelium Reveals Distinct Spatial, Cell Type-Specific, and Temporal Patterns. Immunity 49, 560–575.e6 (2018).

6. Ko, Y. et al. Cell type-specific genes show striking and distinct patterns of spatial expression in the mouse brain. Proc Natl Acad Sci U S A 110, 3095–3100 (2013).

7. Nelson, C. M. Geometric control of tissue morphogenesis. Biochim Biophys Acta 1793, 903–910 (2009).

8. Liu, Q., Hsu, C.-Y. & Shyr, Y. Scalable and model-free detection of spatial patterns and colocalization. Genome Res 32, 1736–1745 (2022).

9. Chen, S. et al. Spatially resolved transcriptomics reveals genes associated with the vulnerability of middle temporal gyrus in Alzheimer’s disease. Acta Neuropathol Commun 10, 188 (2022).

10. Andersson, M. K. et al. The multifunctional FUS, EWS and TAF15 proto-oncoproteins show cell type-specific expression patterns and involvement in cell spreading and stress response. BMC Cell Biol 9, 37 (2008).

11. Maynard, K. R. et al. Transcriptome-scale spatial gene expression in the human dorsolateral prefrontal cortex. Nat Neurosci 24, 425–436 (2021).

12. Rodriques, S. G. et al. Slide-seq: A scalable technology for measuring genome-wide expression at high spatial resolution. Science 363, 1463–1467 (2019).

13. Vickovic, S. et al. High-definition spatial transcriptomics for in situ tissue profiling. Nat Methods 16, 987–990 (2019).

14. Chen, A. et al. Spatiotemporal transcriptomic atlas of mouse organogenesis using DNA nanoball-patterned arrays. Cell 185, 1777–1792.e21 (2022).

15. Cho, C.-S. et al. Microscopic examination of spatial transcriptome using Seq-Scope. Cell 184, 3559–3572.e22 (2021).

16. Stickels, R. R. et al. Highly sensitive spatial transcriptomics at near-cellular resolution with Slide- seqV2. Nat Biotechnol 39, 313–319 (2021).

17. Wang, X. et al. Three-dimensional intact-tissue sequencing of single-cell transcriptional states. Science 361, eaat5691 (2018).

18. Xue, Y. et al. A 3D Atlas of Hematopoietic Stem and Progenitor Cell Expansion by Multi- dimensional RNA-Seq Analysis. Cell Rep 27, 1567–1578.e5 (2019).

19. Zeira, R., Land, M., Strzalkowski, A. & Raphael, B. J. Alignment and integration of spatial transcriptomics data. Nat Methods 19, 567–575 (2022).

20. Young, D. M. et al. Constructing and optimizing 3D atlases from 2D data with application to the developing mouse brain. Elife 10, e61408 (2021).

21. Laurent, J. et al. Convergence of microengineering and cellular self-organization towards functional tissue manufacturing. Nat Biomed Eng 1, 939–956 (2017).

22. Yamada, K. M. & Cukierman, E. Modeling tissue morphogenesis and cancer in 3D. Cell 130, 601– 610 (2007).

23. Lin, J.-R. et al. Multiplexed 3D atlas of state transitions and immune interaction in colorectal cancer. Cell 186, 363–381.e19 (2023).

24. Miller, B. F., Bambah-Mukku, D., Dulac, C., Zhuang, X. & Fan, J. Characterizing spatial gene expression heterogeneity in spatially resolved single-cell transcriptomic data with nonuniform cellular densities. Genome Res. 31, 1843–1855 (2021).

25. Zhu, J., Sun, S. & Zhou, X. SPARK-X: non-parametric modeling enables scalable and robust detection of spatial expression patterns for large spatial transcriptomic studies. Genome Biol 22, 184 (2021).

26. Li, B. et al. Benchmarking spatial and single-cell transcriptomics integration methods for transcript distribution prediction and cell type deconvolution. Nat Methods 19, 662–670 (2022).

27. Hu, J. et al. Statistical and machine learning methods for spatially resolved transcriptomics with histology. Computational and Structural Biotechnology Journal 19, 3829–3841 (2021).

28. Ospina, O. E. et al. Differential gene expression analysis of spatial transcriptomic experiments using spatial mixed models. Sci Rep 14, 10967 (2024).

29. Weber, L. M., Saha, A., Datta, A., Hansen, K. D. & Hicks, S. C. nnSVG for the scalable identification of spatially variable genes using nearest-neighbor Gaussian processes. Nat Commun 14, 4059 (2023).

30. Hao, M., Hua, K. & Zhang, X. SOMDE: a scalable method for identifying spatially variable genes with self-organizing map. Bioinformatics 37, 4392–4398 (2021).

31. Hu, J. et al. SpaGCN: Integrating gene expression, spatial location and histology to identify spatial domains and spatially variable genes by graph convolutional network. Nat Methods 18, 1342–1351 (2021).

32. Wang, J. et al. Dimension-agnostic and granularity-based spatially variable gene identification using BSP. Nat Commun 14, 7367 (2023).

33. Arya, S. An Optimal Algorithm for Approximate Nearest Neighbor Searching in Fixed Dimensions.

34. Svensson, V., Teichmann, S. A. & Stegle, O. SpatialDE: identification of spatially variable genes. Nat Methods 15, 343–346 (2018).

35. Sun, S., Zhu, J. & Zhou, X. Statistical analysis of spatial expression patterns for spatially resolved transcriptomic studies. Nat Methods 17, 193–200 (2020).

36. Tian, T., Zhang, J., Lin, X., Wei, Z. & Hakonarson, H. Dependency-aware deep generative models for multitasking analysis of spatial omics data. Nat Methods 21, 1501–1513 (2024).

37. Jiang, F. et al. Simultaneous profiling of spatial gene expression and chromatin accessibility during mouse brain development. Nat Methods 20, 1048–1057 (2023).

38. Zhang, D. et al. Spatial epigenome–transcriptome co-profiling of mammalian tissues. Nature 616, 113–122 (2023).

39. Machanick, P. & Bailey, T. L. MEME-ChIP: motif analysis of large DNA datasets. Bioinformatics 27, 1696–1697 (2011).

40. Dolfini, D., Imbriano, C. & Mantovani, R. The role(s) of NF-Y in development and differentiation. Cell Death Differ 1–12 (2024) doi:10.1038/s41418-024-01388-1.

41. Tsuchiya, Y. et al. The Casein Kinase 2-Nrf1 Axis Controls the Clearance of Ubiquitinated Proteins by Regulating Proteasome Gene Expression. Mol Cell Biol 33, 3461–3472 (2013).

42. Harvey, J. D. & Heinbockel, T. Neuromodulation of Synaptic Transmission in the Main Olfactory Bulb. Int J Environ Res Public Health 15, 2194 (2018).

43. Pérez-Revuelta, L. et al. Secretagogin expression in the mouse olfactory bulb under sensory impairments. Sci Rep 10, 21533 (2020).

44. Lledo, P.-M. & Valley, M. Adult Olfactory Bulb Neurogenesis. Cold Spring Harb Perspect Biol 8, a018945 (2016).

45. Li, S. et al. Asynchronous release sites align with NMDA receptors in mouse hippocampal synapses. Nat Commun 12, 677 (2021).

46. Han, K. A. et al. Specification of neural circuit architecture shaped by context-dependent patterned LAR-RPTP microexons. Nat Commun 15, 1624 (2024).

47. Ferreira, R. M. et al. A MEF2C transcription factor network regulates proliferation of glomerular endothelial cells in diabetic kidney disease. 2024.09.27.615250 Preprint at 10.1101/2024.09.27.615250 (2024).

48. Lake, B. B. et al. An atlas of healthy and injured cell states and niches in the human kidney. Nature 619, 585–594 (2023).

49. Sarakpi, T., Mesic, A. & Speer, T. Leukocyte–endothelial interaction in CKD. Clinical Kidney Journal 16, 1845–1860 (2023).

50. Wolf, F. A., Angerer, P. & Theis, F. J. SCANPY: large-scale single-cell gene expression data analysis. Genome Biology 19, 15 (2018).

51. Bentley, J. L. Multidimensional binary search trees used for associative searching. Commun. ACM 18, 509–517 (1975).

52. Svensson, V., Teichmann, S. A. & Stegle, O. SpatialDE: identification of spatially variable genes. Nat Methods 15, 343–346 (2018).

53. Sun, S., Zhu, J. & Zhou, X. Statistical analysis of spatial expression patterns for spatially resolved transcriptomic studies. Nat Methods 17, 193–200 (2020).

54. Li, Z. et al. Benchmarking computational methods to identify spatially variable genes and peaks. bioRxiv 2023.12.02.569717 (2023) doi:10.1101/2023.12.02.569717.

55. Chen, C., Kim, H. J. & Yang, P. Evaluating spatially variable gene detection methods for spatial transcriptomics data. Genome Biology 25, 18 (2024).

56. Janesick, A. et al. High resolution mapping of the tumor microenvironment using integrated single-cell, spatial and in situ analysis. Nat Commun 14, 8353 (2023).

57. Wu, T. et al. clusterProfiler 4.0: A universal enrichment tool for interpreting omics data. Innovation 2, (2021).

