## Supplementary File for "scBSP: A fast and accurate tool for identifying spatially variable features from high-resolution spatial omics data"

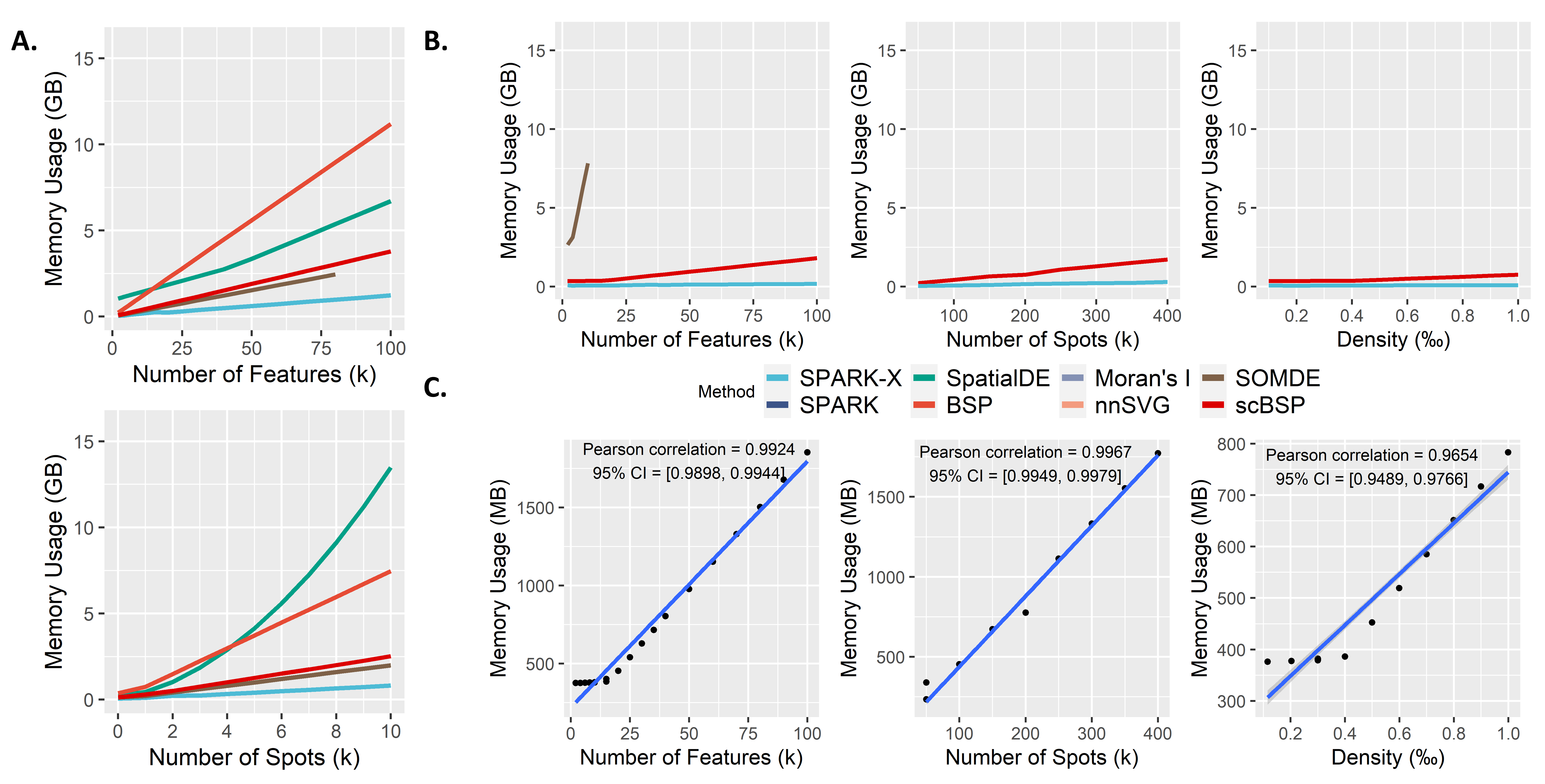

Supplementary Figure 1. A: Memory usage (y-axis) for analyzing spatial omics data comprising 20,000 features across 3,000 spots. B: Memory usage (y-axis) for analyzing high-resolution spatial omics data comprising 20,000 features across 100,000 spots, with a data density of 0.0005. This analysis varies one parameter while keeping the other two constants. C: Memory usage (y-axis) of scBSP on the high-resolution spatial omics data (run n = 10 times on a single processor core) with varied number of features, spot count, and data density (x-axis).

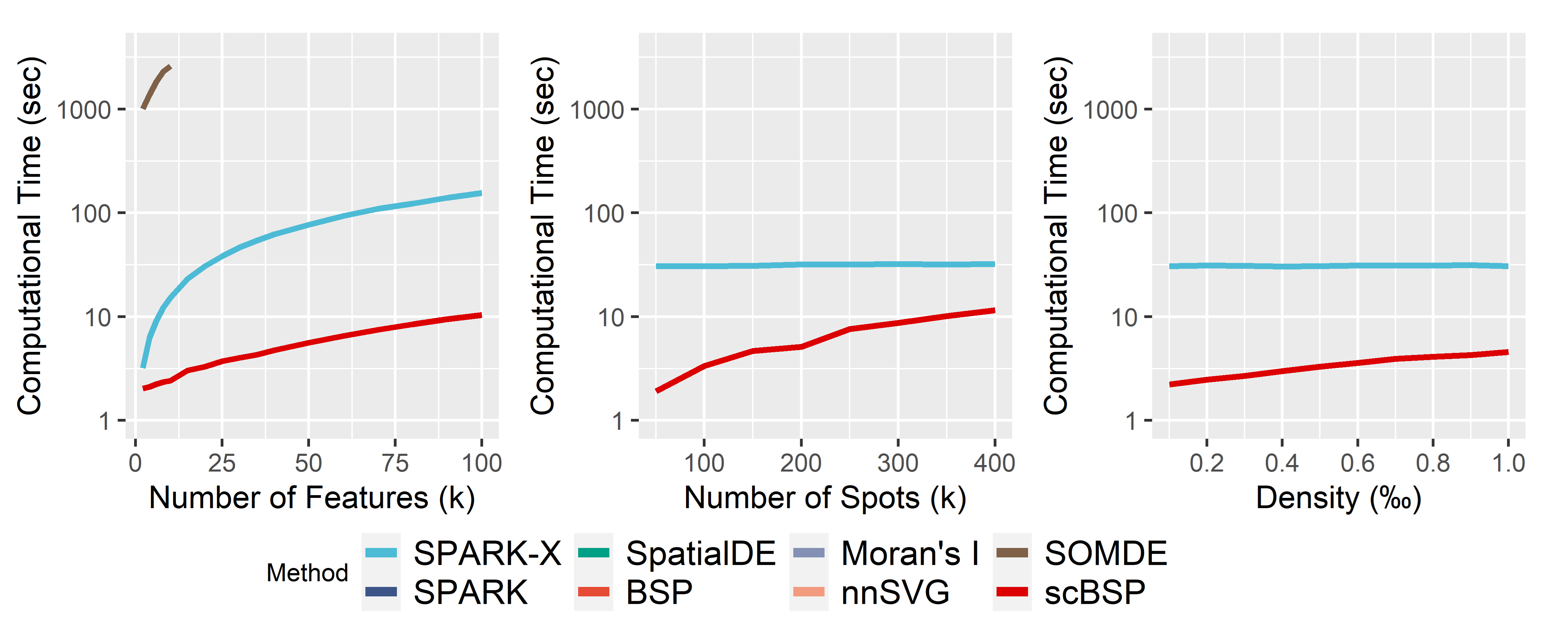

Supplementary Figure 2. Computational time (y-axis) for analyzing high-resolution spatial omics data comprising 20,000 genes across 100,000 spots, with a data density of 0.0005. This analysis varies one parameter while keeping the other two constants.

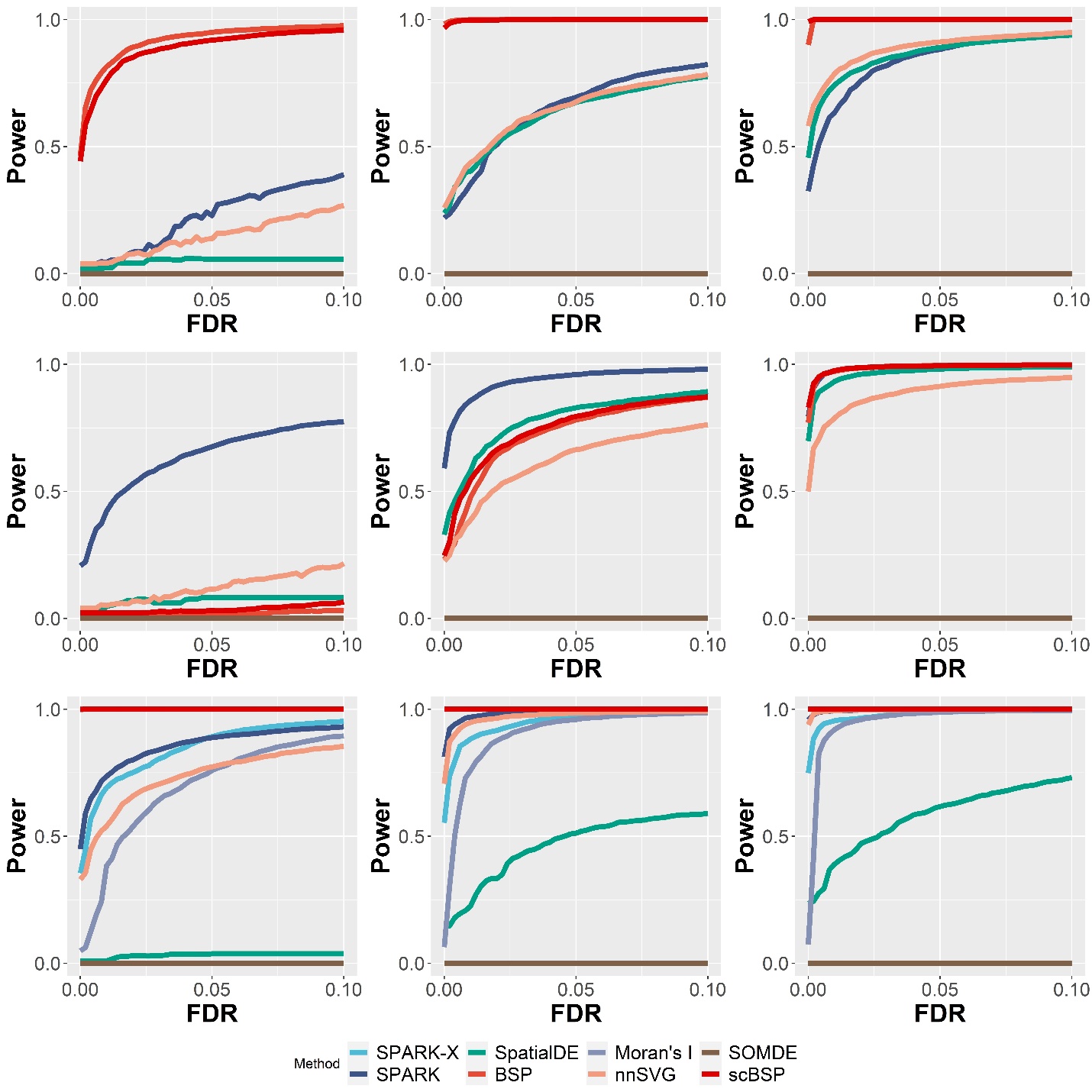

Supplementary Figure 3. Statistical power on 2D simulations with varied signal strengths. Signal strengths were measured as the fold changes in the averaged expressions between the pattern and non-pattern regions. Power curves were drawn using the averaged statistical power (y-axis) across ten replicates against the false discovery rates (x-axis) for the detected SVGs from each method. Results with weak (FC = 3), moderate (FC = 4), and high (FC = 5) signal strengths are shown in the left, middle, and right columns, while the upper, middle, and bottom rows represent the results from three spatial patterns in Fig.2 A. All simulations were generated using a fixed moderate noise level.

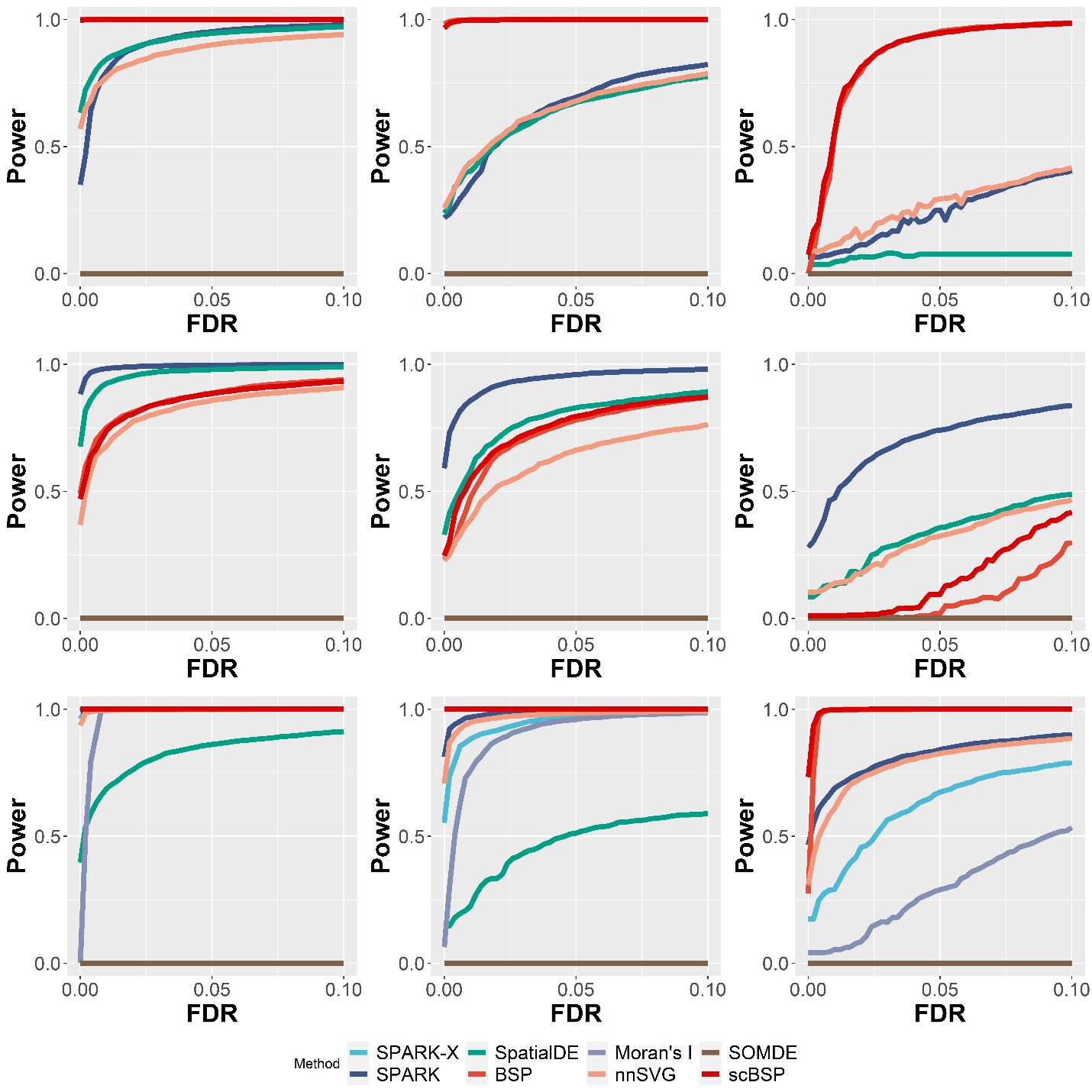

Supplementary Figure 4. Statistical power on 2D simulations with varied noise levels. Noise levels were defined as the dispersion parameters ($\tau_{2}$) in SPARK’s model. Power curves were drawn using the averaged statistical power (y-axis) across ten replicates against the false discovery rates (x-axis) for the detected SVGs from each method. Results with low ($\tau_{2}$=0.2), moderate ($\tau_{2}$=0.5), and high ($\tau_{2}$=0.8) noise levels are shown in the left, middle, and right columns, while the upper, middle, and bottom rows represent the results from three spatial patterns in Fig.2 A. All simulations were generated using a fixed moderate signal strength.

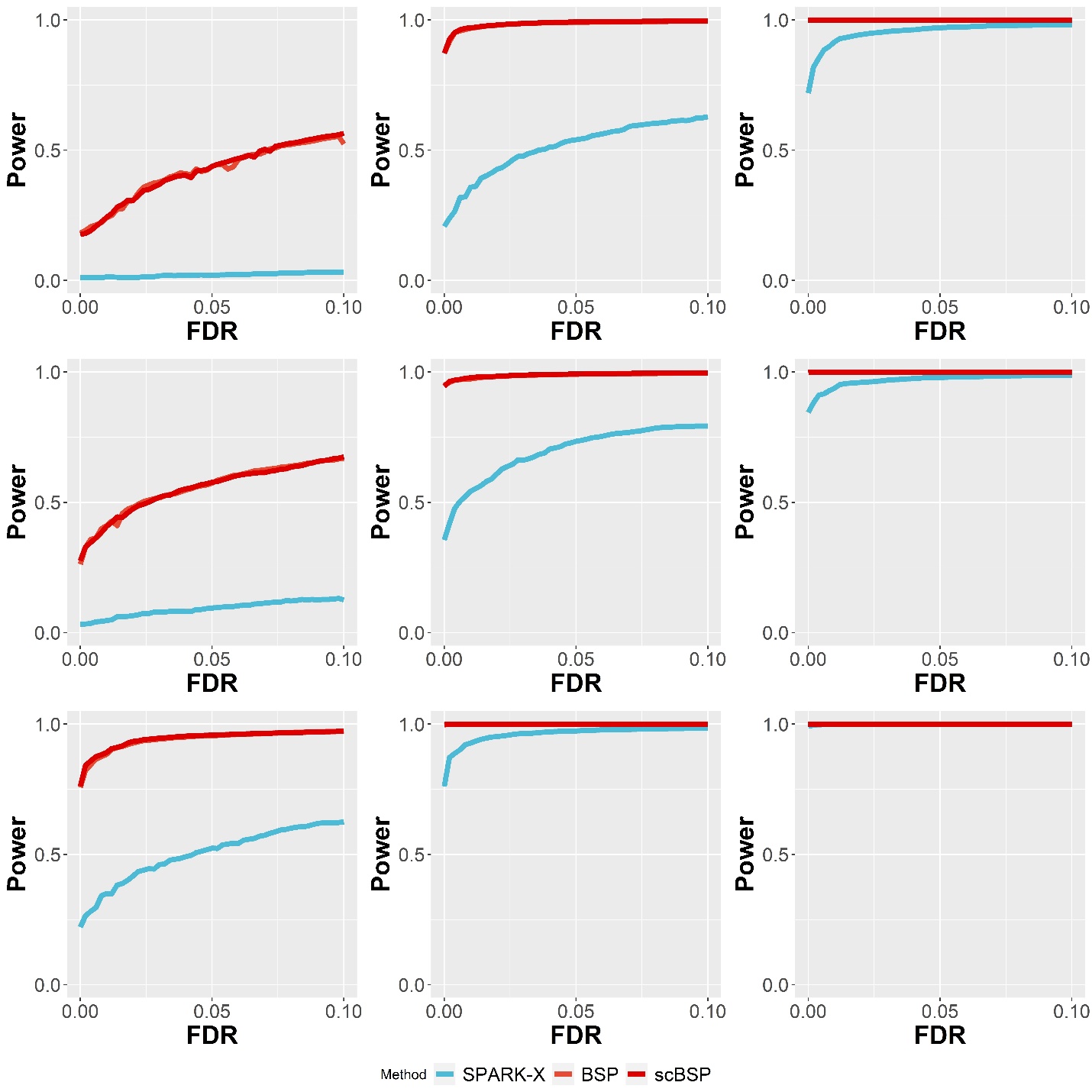

Supplementary Figure 5. Statistical power on 3D simulations of continuous patterns with varied pattern sizes. Pattern size was measured as the radius of the pattern as described in the Method section. Power curves were drawn using the averaged statistical power (y-axis) across ten replicates against the false discovery rates (x-axis) for the detected SVGs from each method. Results with small (radius=1.5), moderate (radius=2.0), and large (radius=2.5) pattern sizes are shown in the left, middle, and right columns, while the upper, middle, and bottom rows represent the results from three continuous spatial patterns in Fig.2 C. All simulations were generated using a fixed moderate signal strength and noise level.

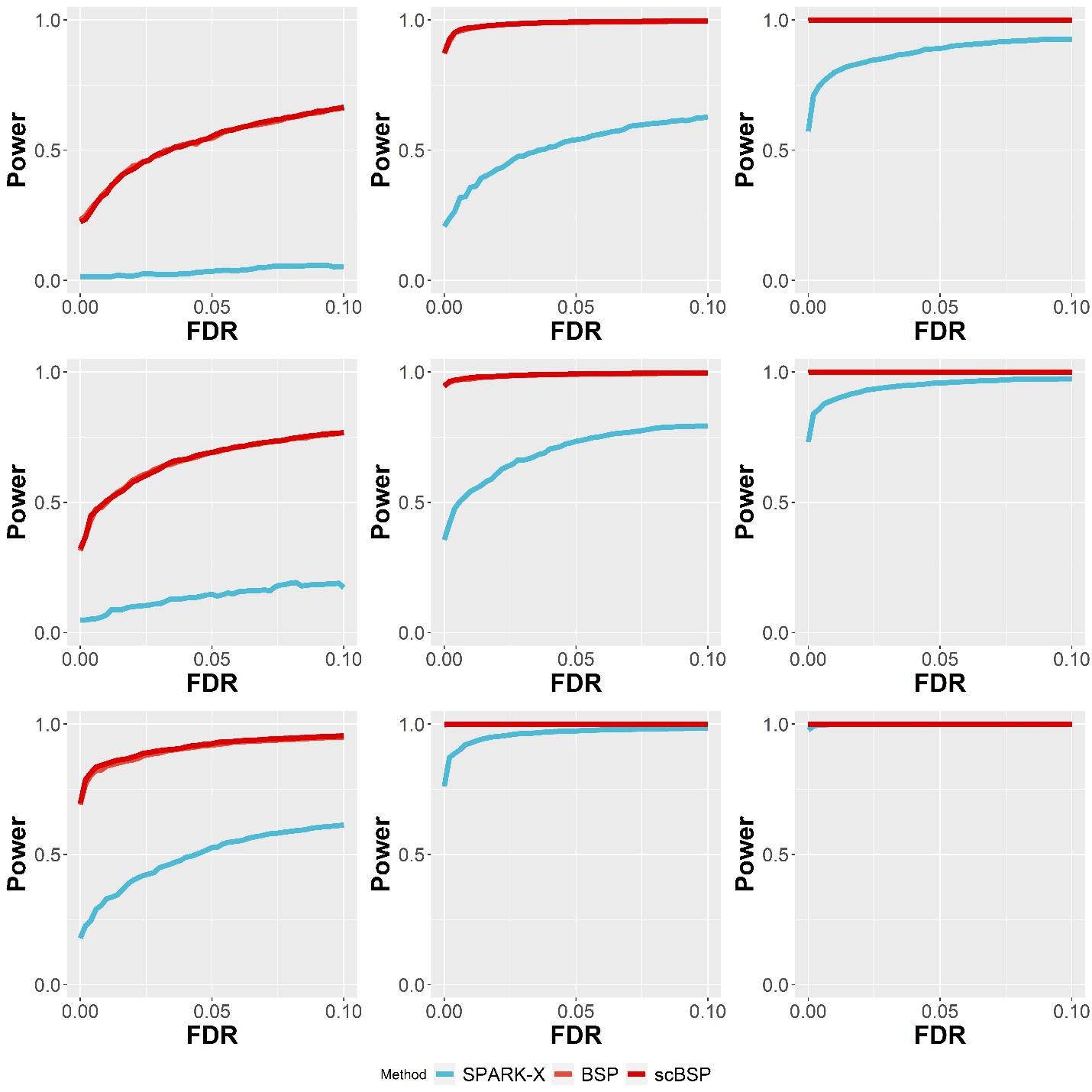

Supplementary Figure 6. Statistical power on 3D simulations of continuous patterns with varied signal strengths. Signal strengths were measured as the fold changes in the averaged expressions between the pattern and non-pattern regions. Power curves were drawn using the averaged statistical power (y-axis) across ten replicates against the false discovery rates (x-axis) for the detected SVGs from each method. Results with weak (FC = 2.0), moderate (FC = 2.5), and high (FC = 3.0) signal strengths are shown in the left, middle, and right columns, while the upper, middle, and bottom rows represent the results from three continuous spatial patterns in Fig.2 C. All simulations were generated using a fixed moderate pattern size and noise level.

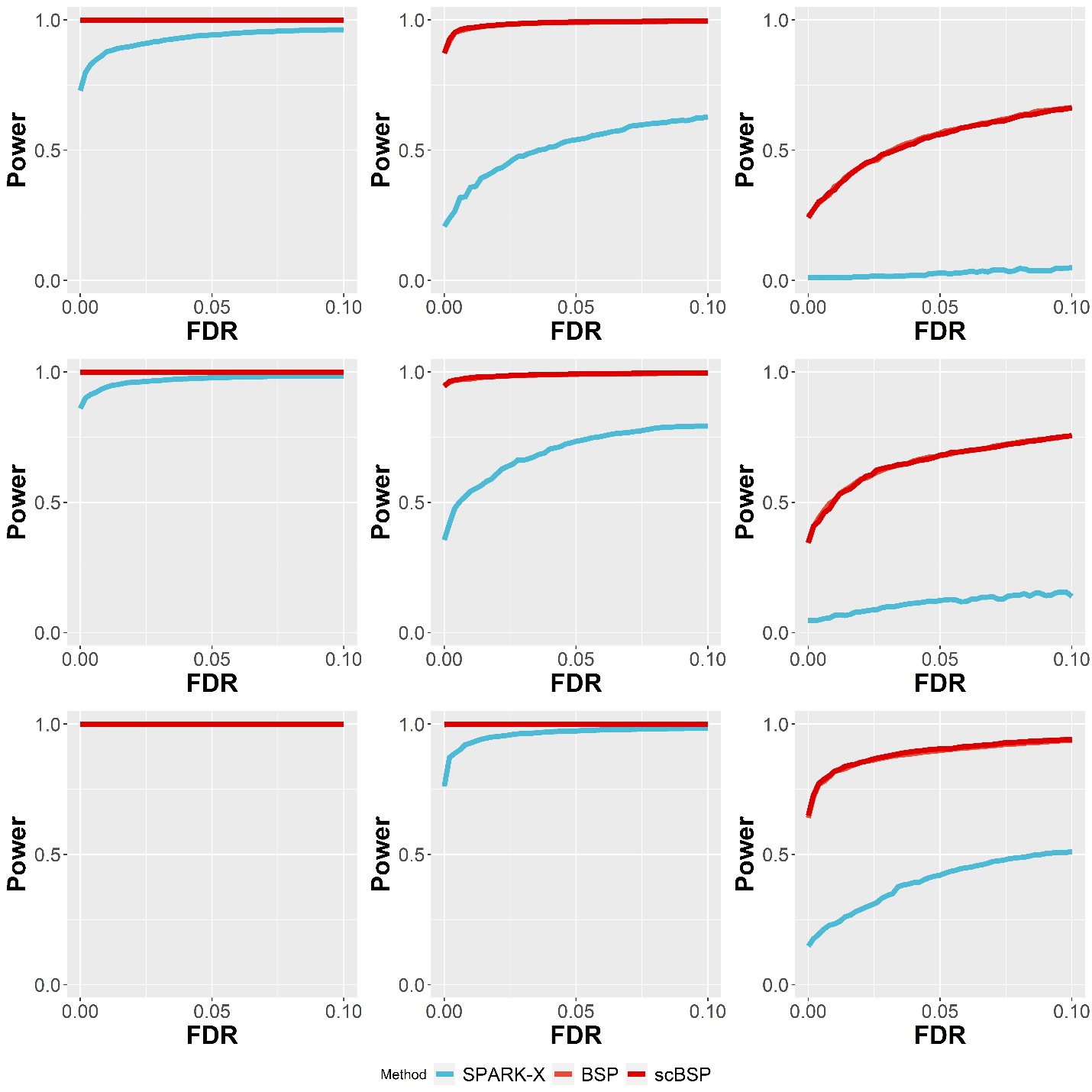

Supplementary Figure 7. Statistical power on 3D simulations of continuous patterns with varied noise levels. Noise levels ($\tau$) were measured as the proportions to the averaged standard deviation of simulated genes (detailed in the Method section). Power curves were drawn using the averaged statistical power (y-axis) across ten replicates against the false discovery rates (x-axis) for the detected SVGs from each method. Results with low ($\tau=0$), moderate ($\tau=1$), and high ($\tau=2$) noise levels are shown in the left, middle, and right columns, while the upper, middle, and bottom rows represent the results from three continuous spatial patterns in Fig.2 C. All simulations were generated using a fixed moderate pattern size and signal strength.

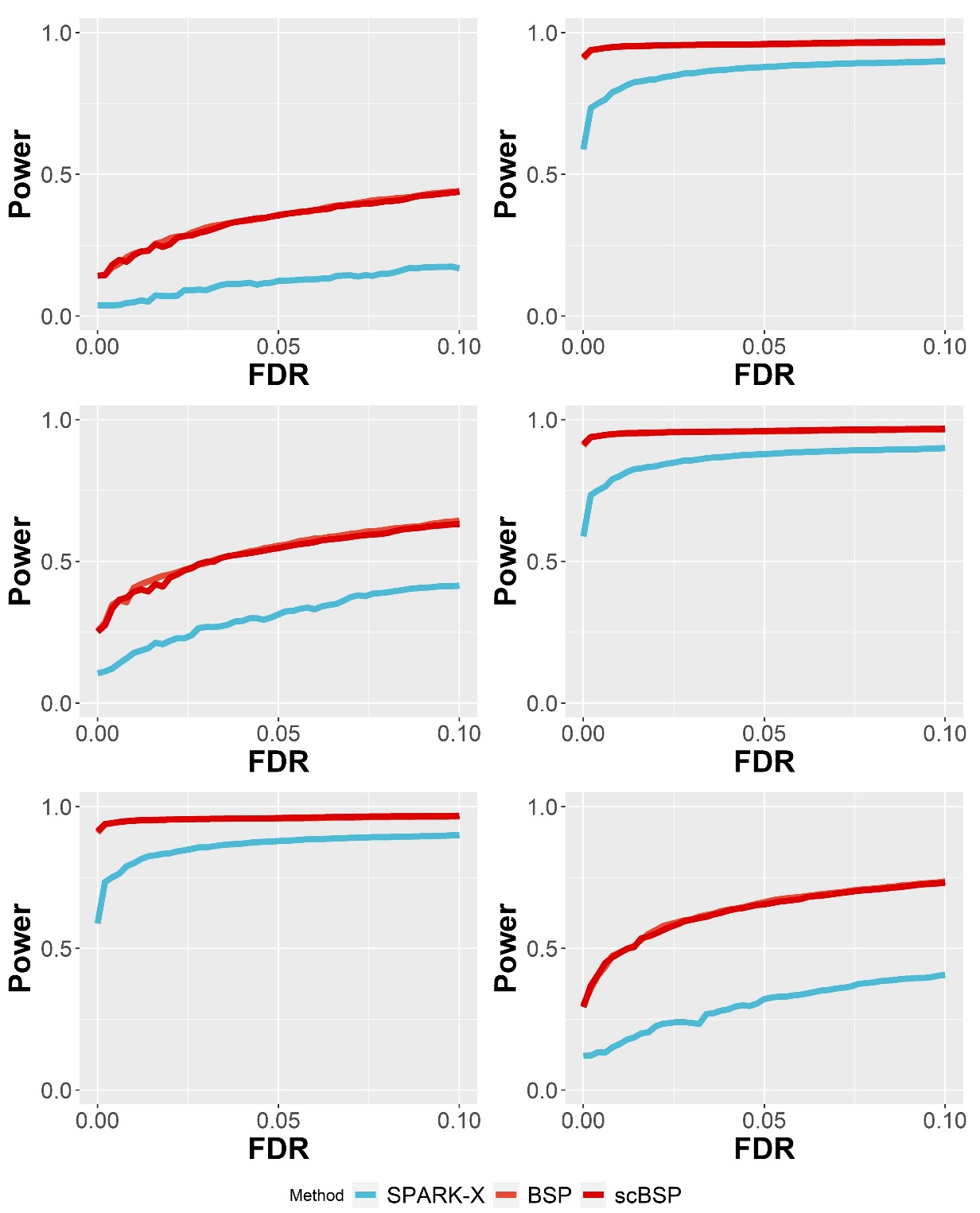

Supplementary Figure 8. Statistical power on 3D simulations of discrete patterns. Power curves were drawn using the averaged statistical power (y-axis) across ten replicates against the false discovery rates (x-axis) for the detected SVGs from each method. Results with varied pattern size (left: radius=1.5; right: radius=2.0), signal strengths (left: FC=2.0; right: FC=2.5) and noise levels (left: $\tau=2.0$; right: $\tau=3.0$) are shown in the upper, middle, and bottom rows as described in the Method section.

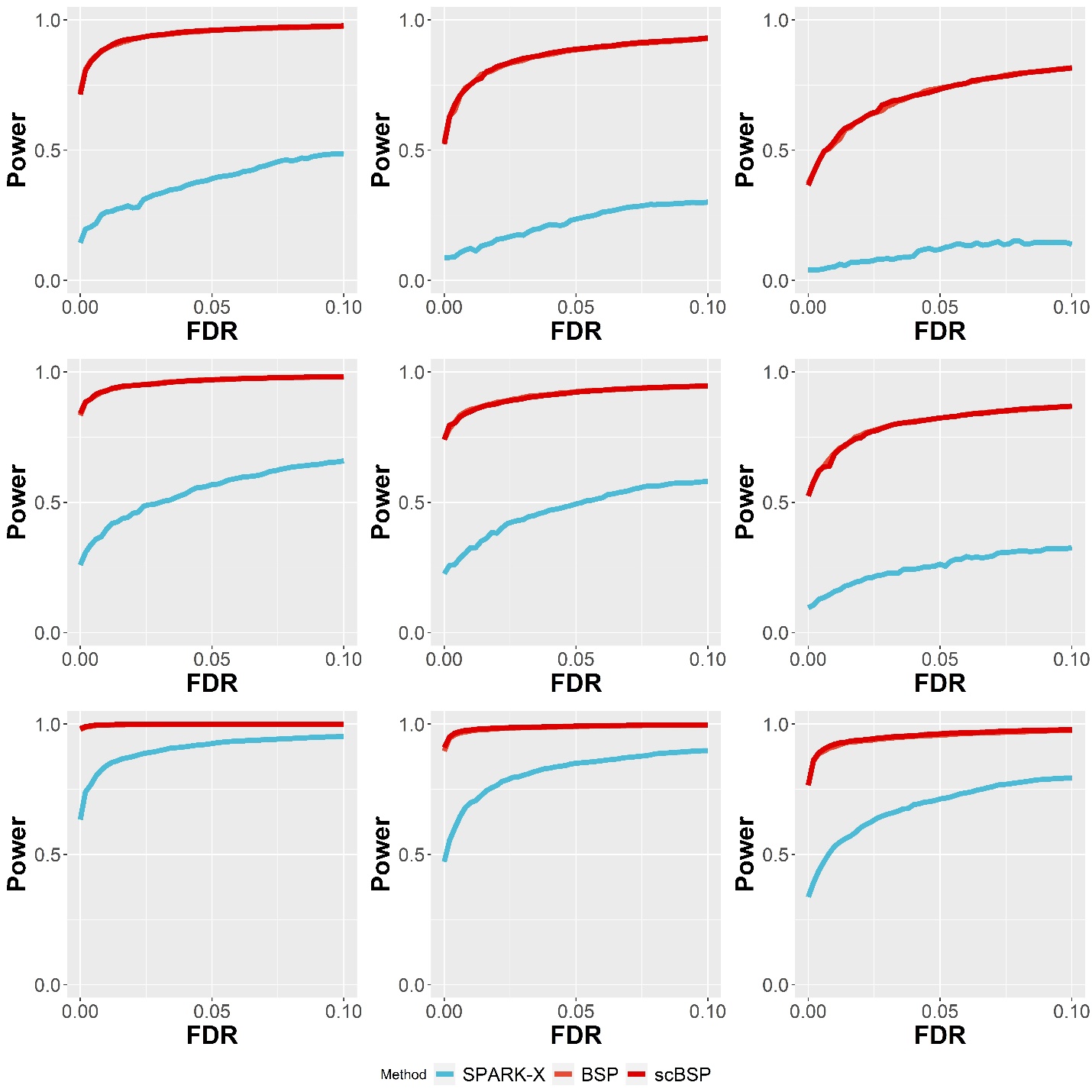

Supplementary Figure 9. Statistical power on 3D simulations of continuous patterns with varied dropout rates. Power curves were drawn using the averaged statistical power (y-axis) across ten replicates against the false discovery rates (x-axis) for the detected SVGs from each method. Results with low (10%), moderate (20%), and high (30%) dropout rates are shown in the left, middle, and right columns, while the upper, middle, and bottom rows represent the results from three continuous spatial patterns in Fig.2 C. All simulations were generated using a fixed moderate pattern size and signal strength.

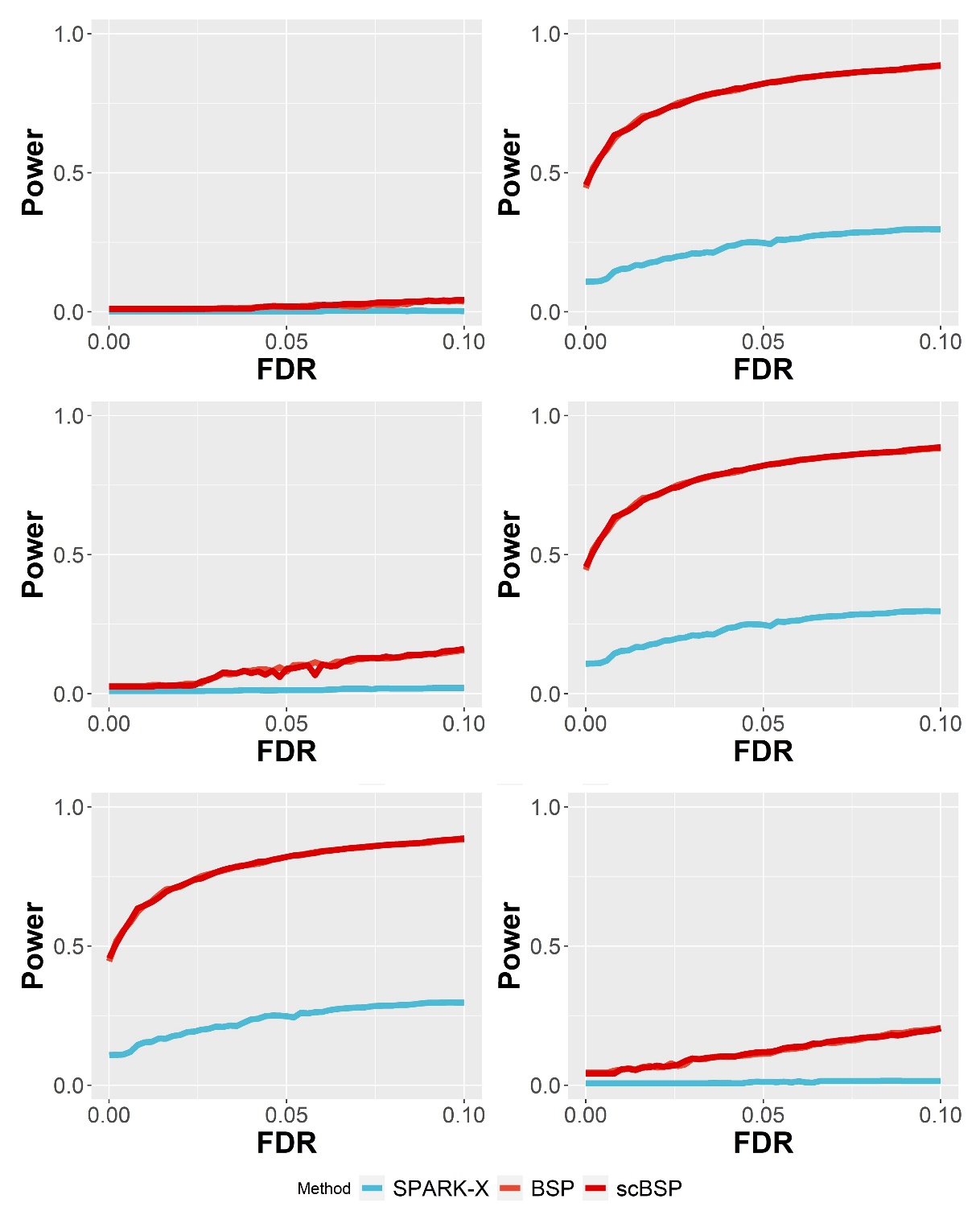

Supplementary Figure 10. Statistical power on 3D simulations of with inconsistent inter-plane and within-plane spatial resolution. Power curves were drawn using the averaged statistical power (y-axis) across ten replicates against the false discovery rates (x-axis) for the detected SVGs from each method. Results with varied pattern size (left: radius=1.5; right: radius=2.0), signal strengths (left: FC=2.0; right: FC=2.5) and noise levels (left: $\tau=1.0$; right: $\tau=2.0$) are shown in the upper, middle, and bottom rows as described in the Method section.

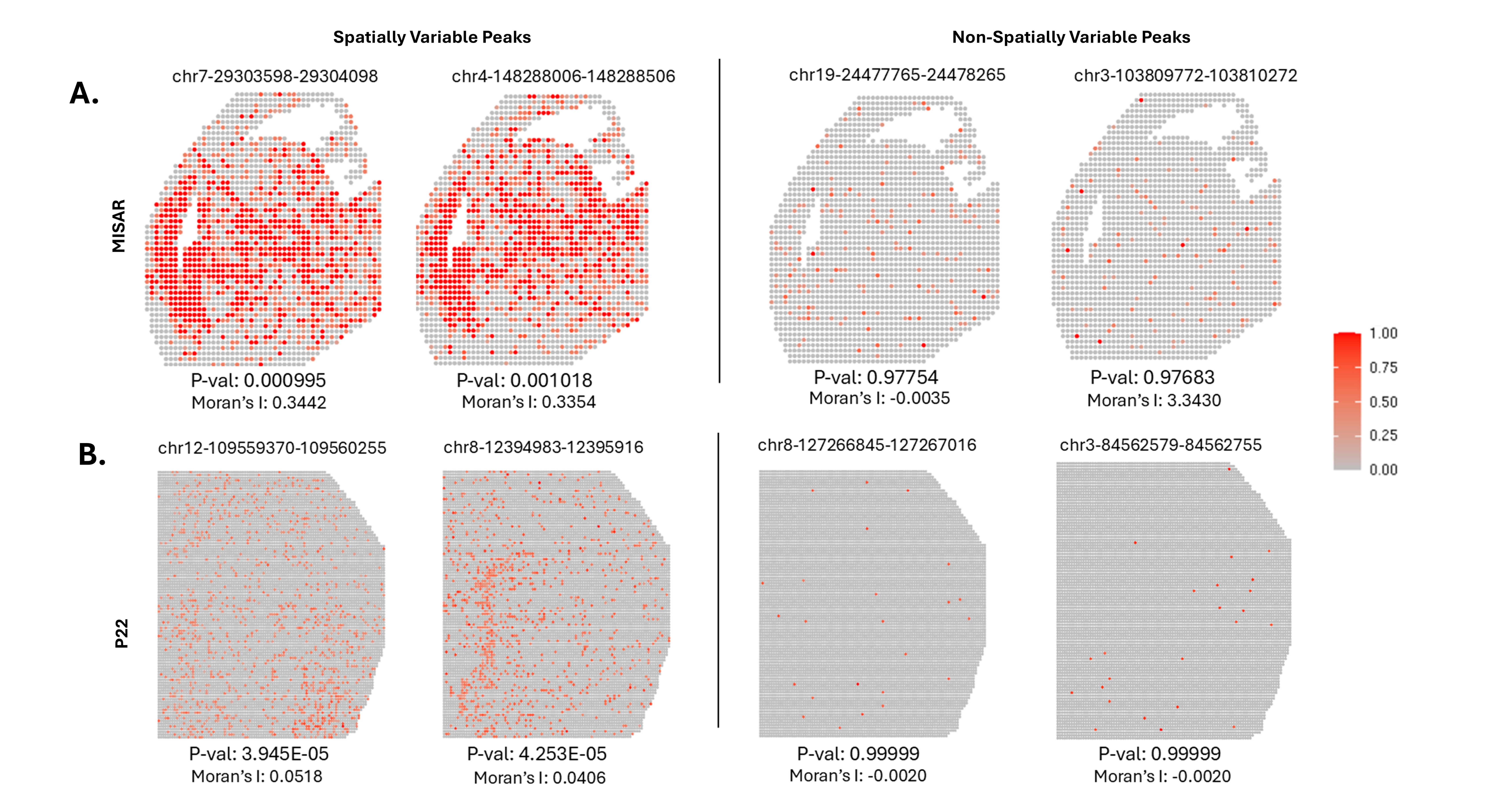

Supplementary Figure 11. Additional top significant and insignificant spatially variable peaks identified by scBSP. A: Additional top significant and insignificant spatially variable peaks on MISAR mouse brain dataset. B: Additional top significant and insignificant spatially variable peaks on the P22 mouse brain dataset.

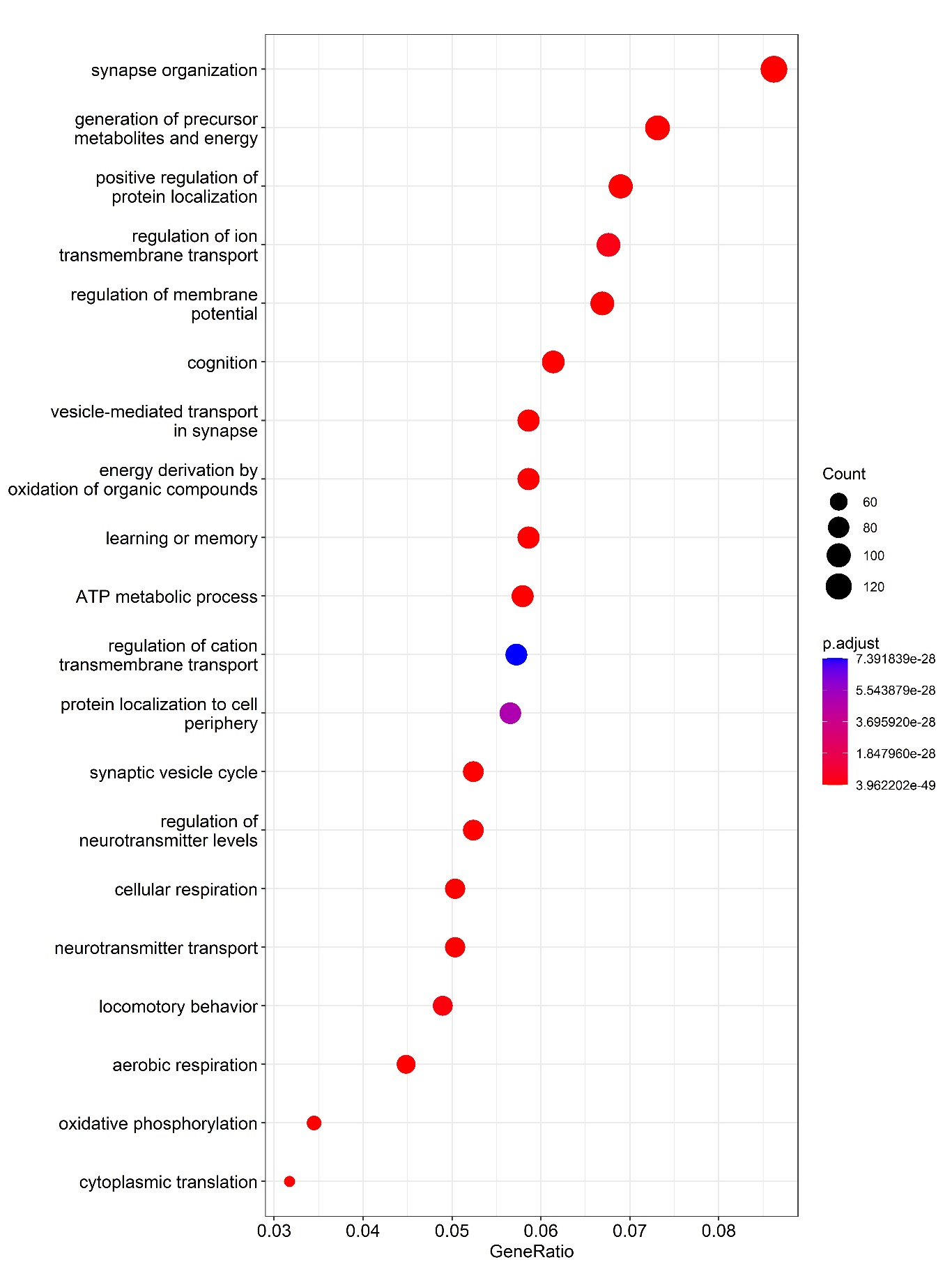

Supplementary Figure 12. Enriched gene ontology terms on 10x Visium mouse brain anterior 1 data.

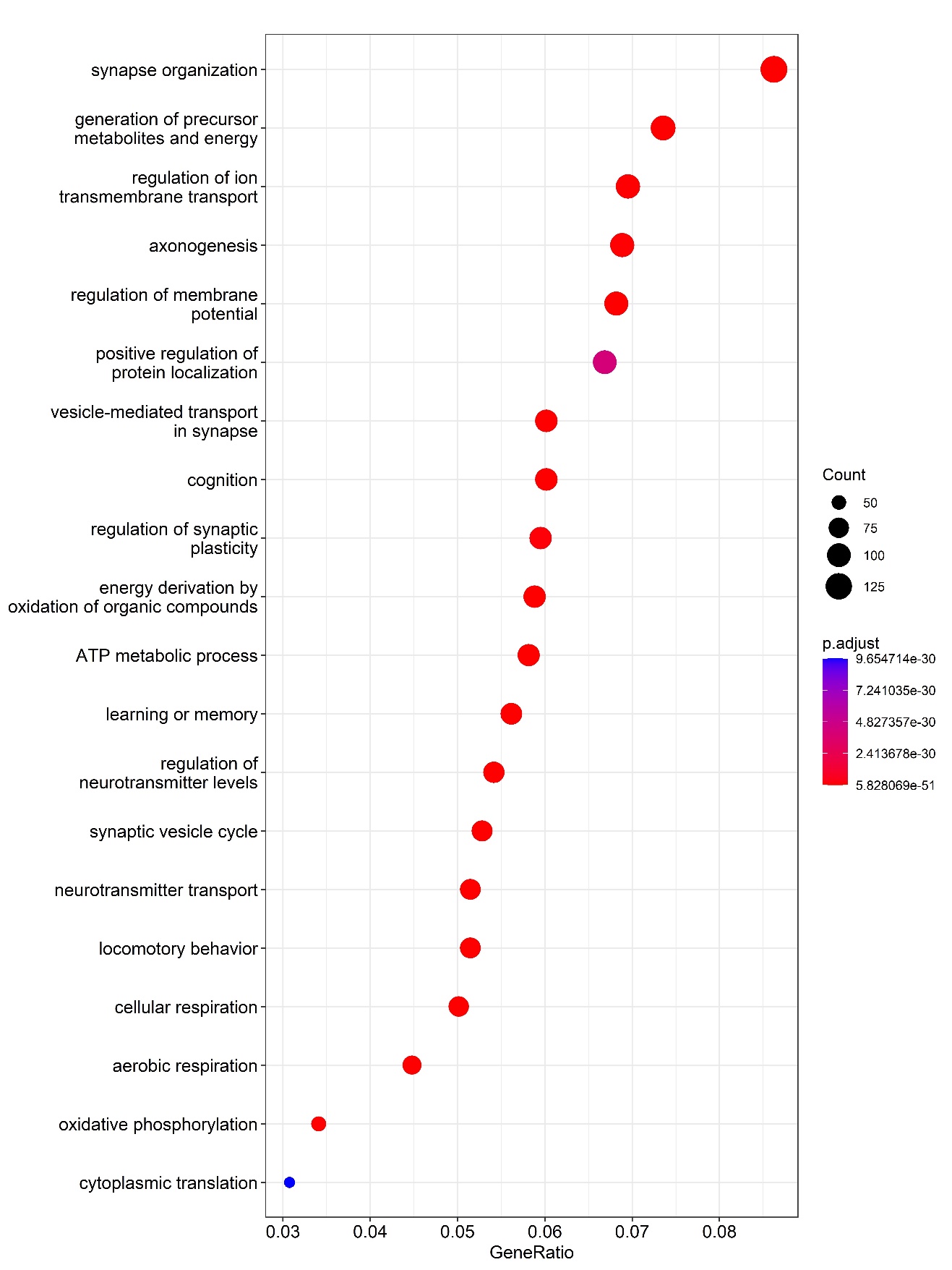

Supplementary Figure 13. Enriched gene ontology terms on 10x Visium mouse brain anterior 2 data.

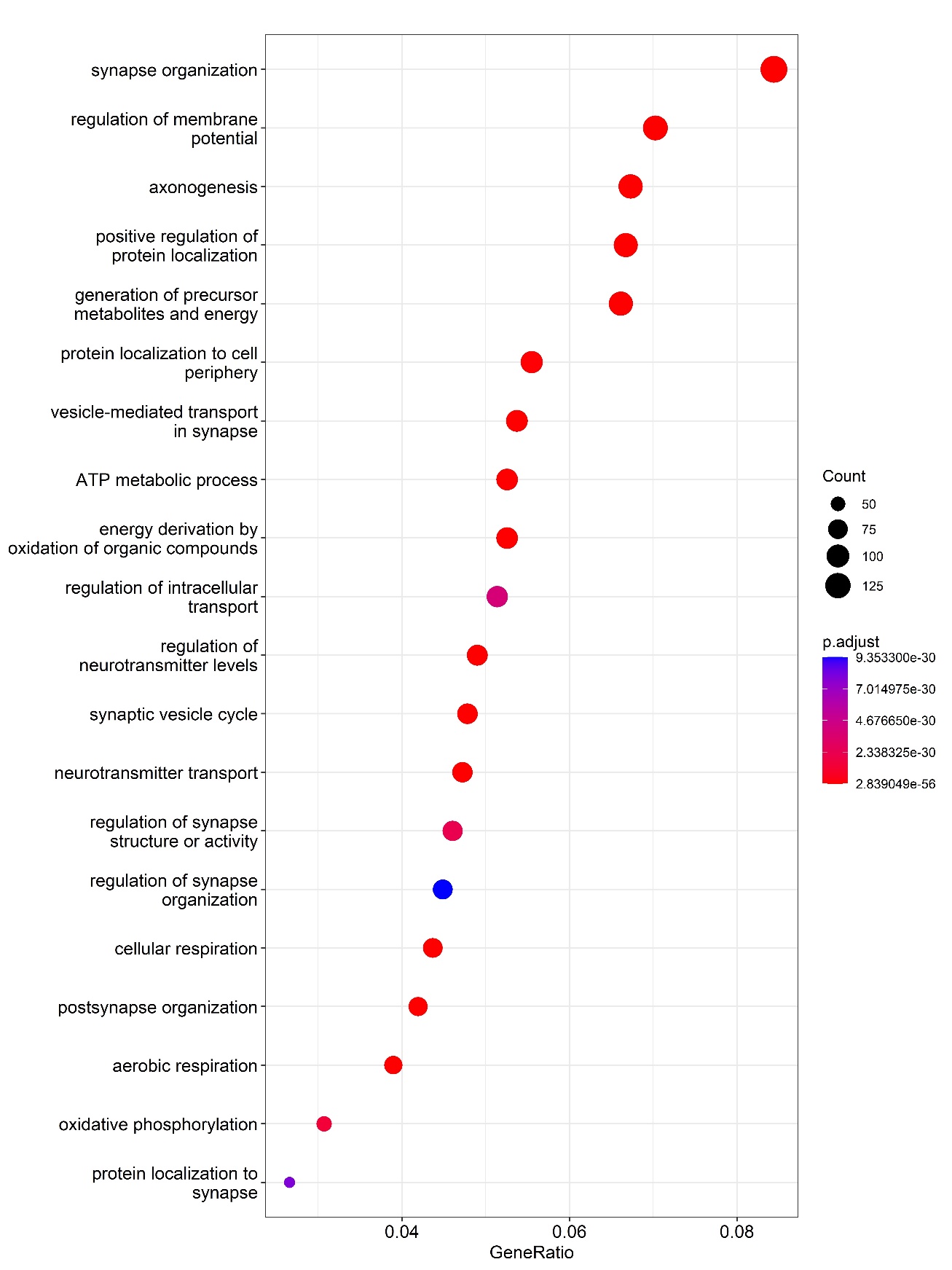

Supplementary Figure 14. Enriched gene ontology terms on 10x Visium mouse brain Posterior 1 data.

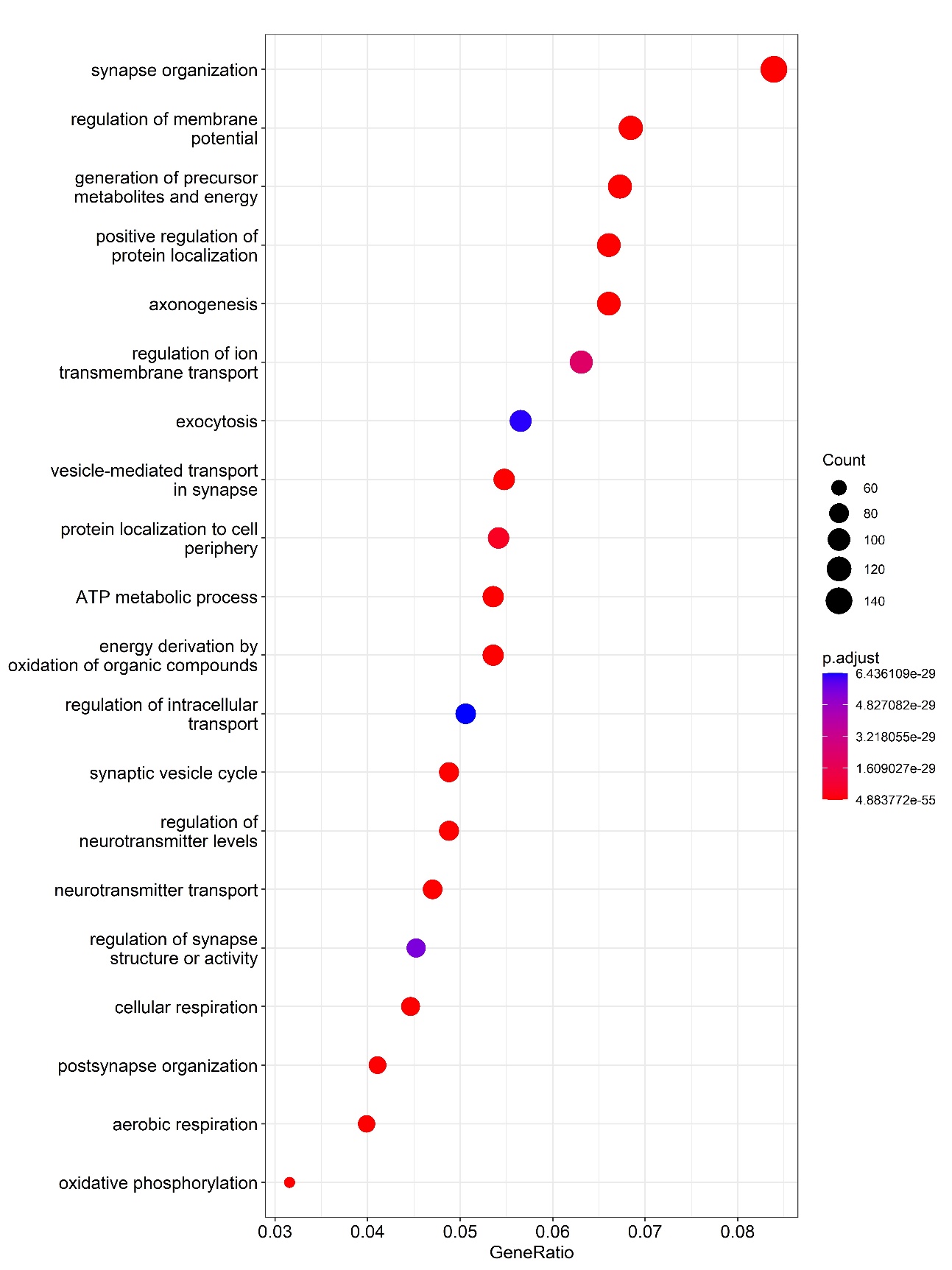

Supplementary Figure 15. Enriched gene ontology terms on 10x Visium mouse brain Posterior 2 data.

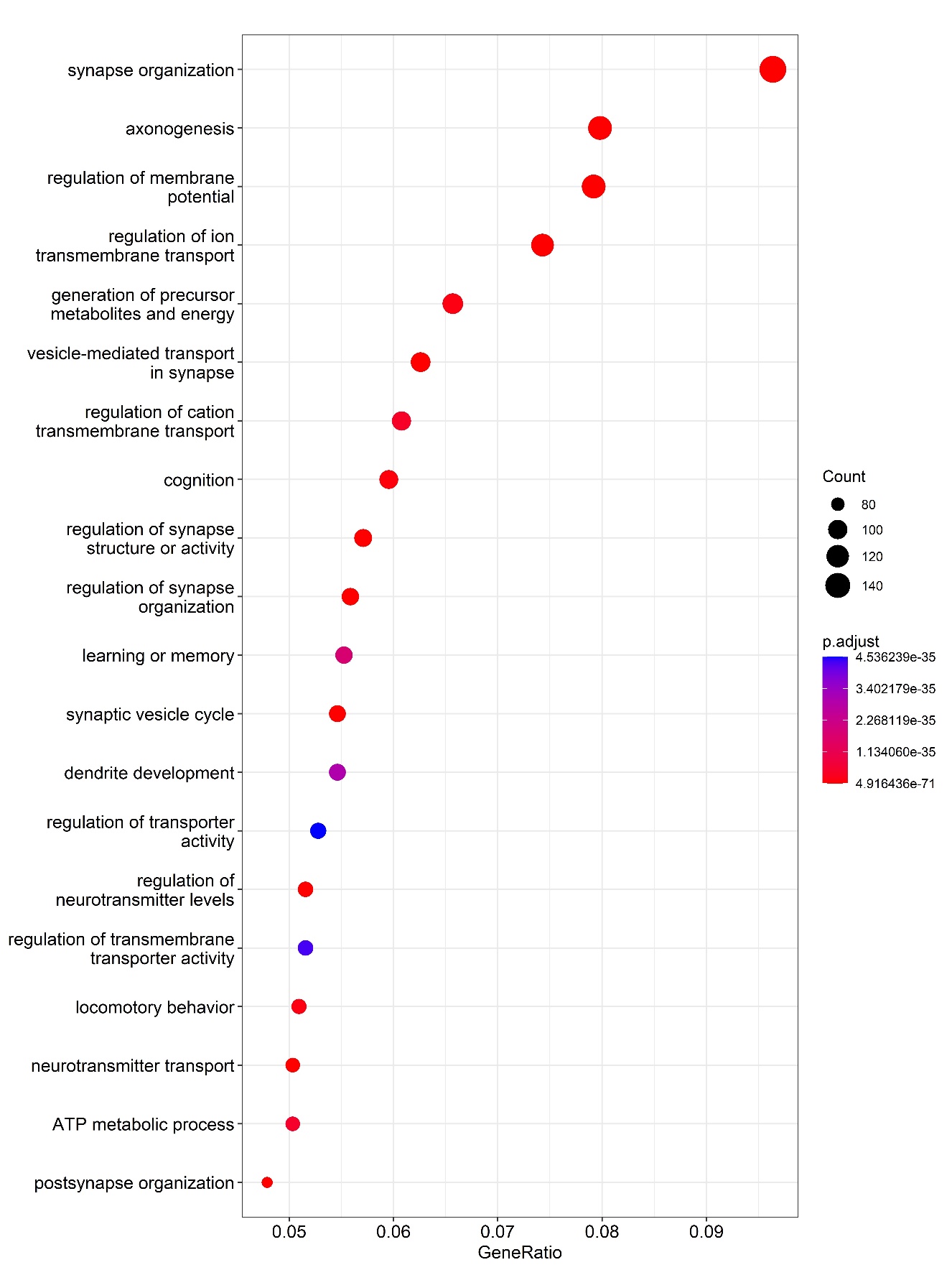

Supplementary Figure 16. Enriched gene ontology terms on Stereo-seq mouse brain data.

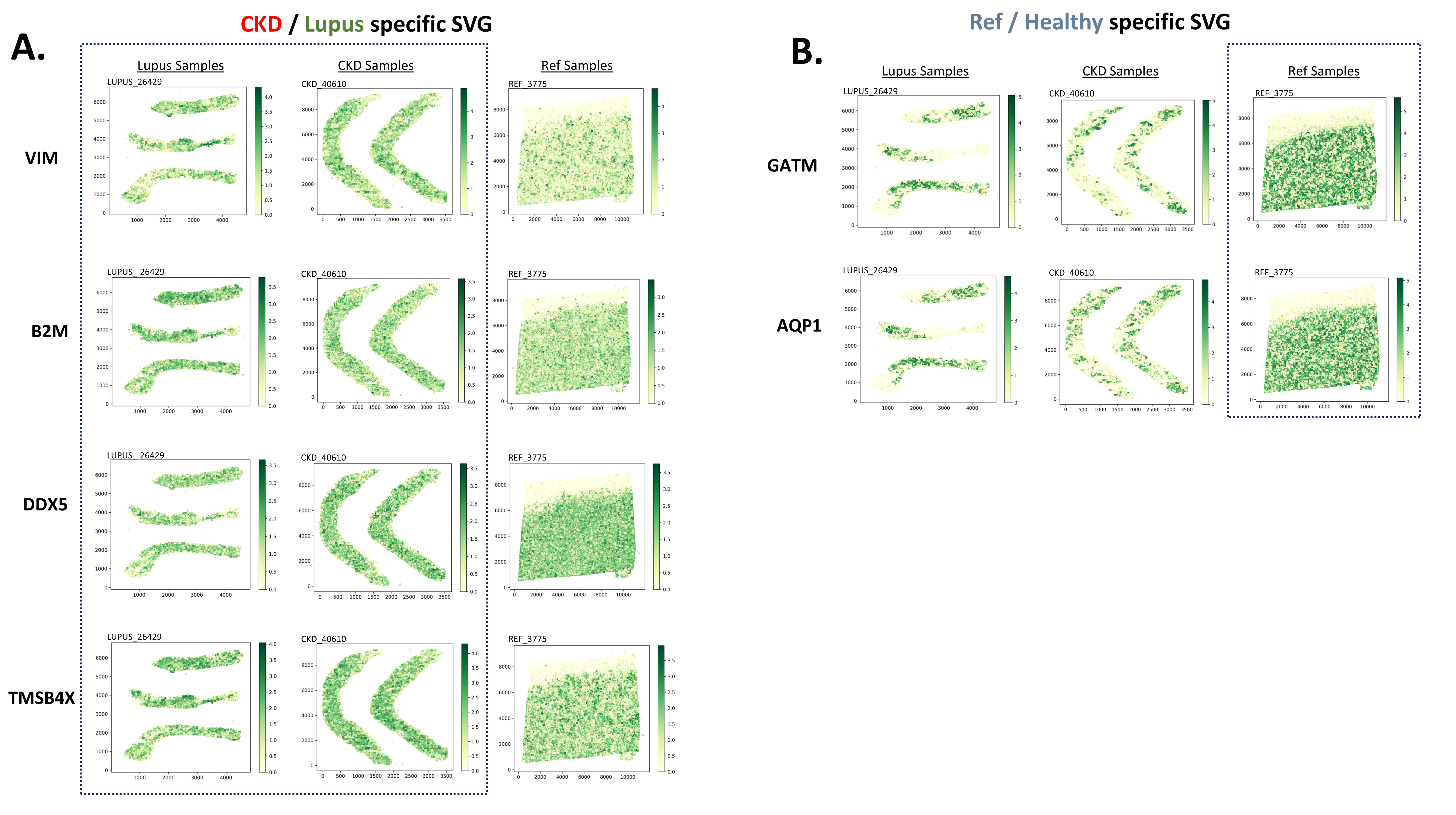

Supplementary Figure 17. Gene expression plots of the 10x Xenium kidney samples. **A**. Plots the gene expression of CKD/Lupus specific SVGs VIM, B2M, DDX5, and TMSB4X **B**. Plots the gene expression of reference specific SVGs GATM and AQP1. The difference in expression is observed in the CKD/Lupus vs. reference samples.

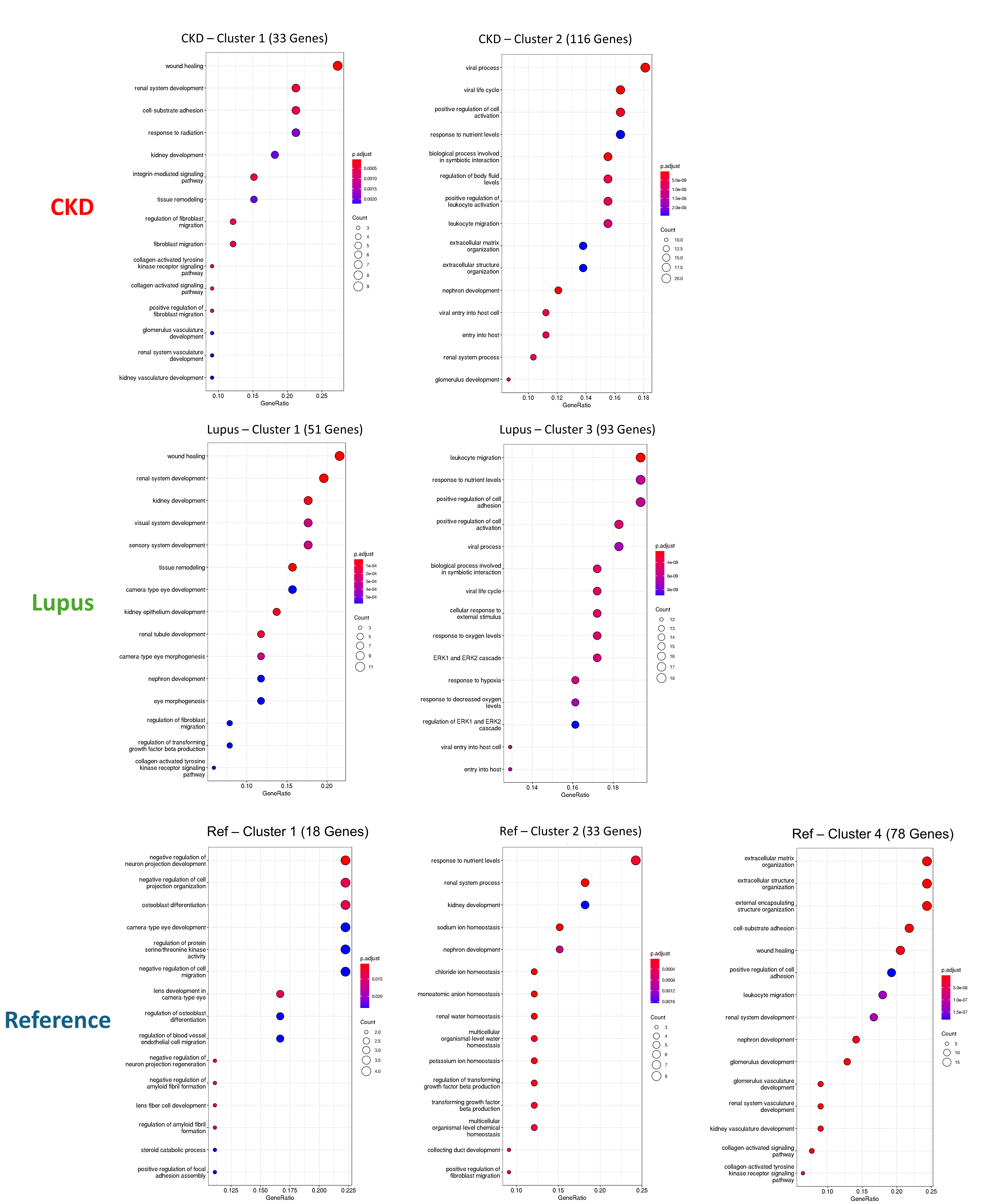

Supplementary Figure 18. Gene Ontology terms comparison between different condition clusters, with clusters of < 10 genes omitted.

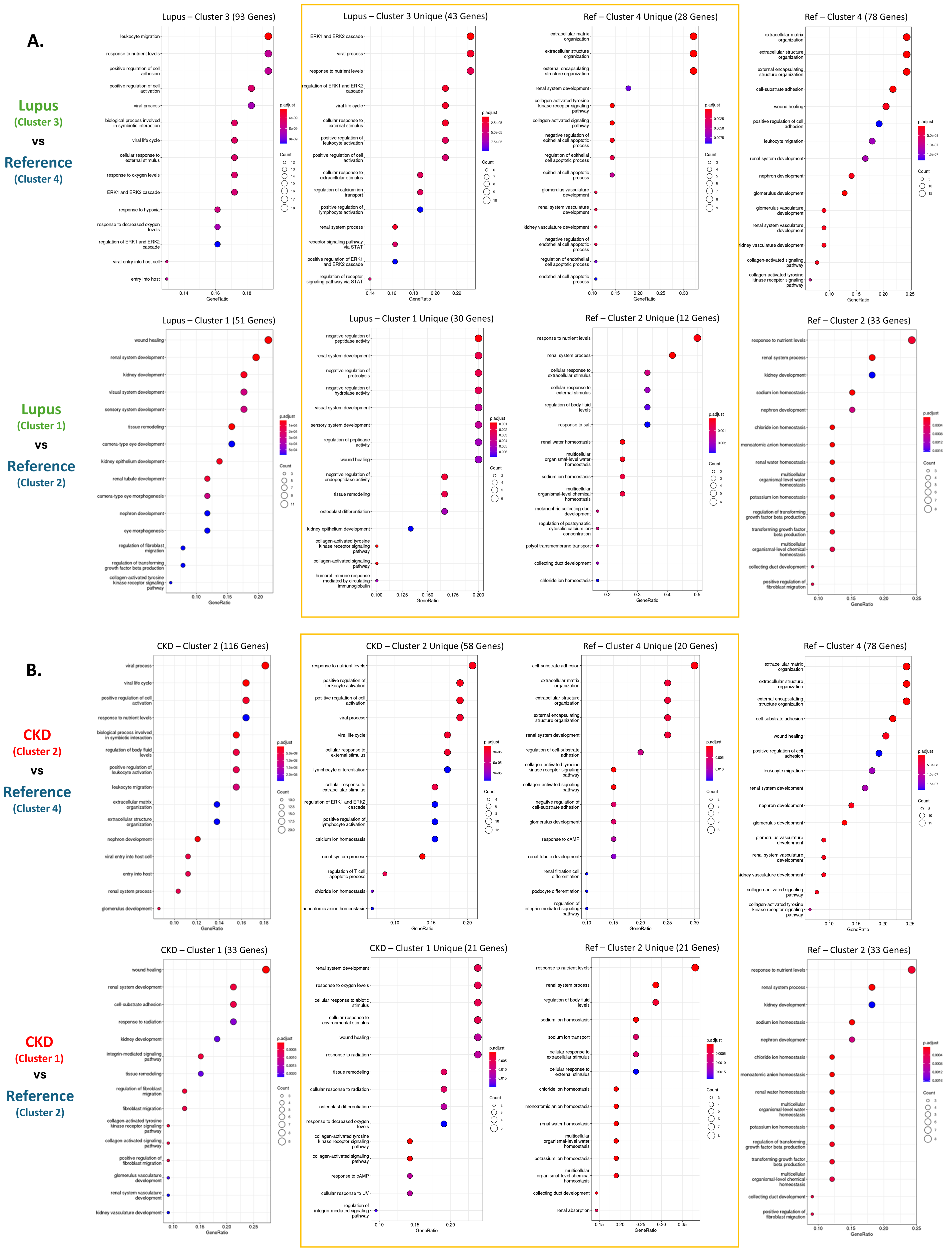

Supplementary Figure 19. Gene Ontology terms comparison between full clusters and unique genes within the cluster. **A.** Lupus vs. Reference condition comparisons **B.** CKD vs. Reference comparisons.

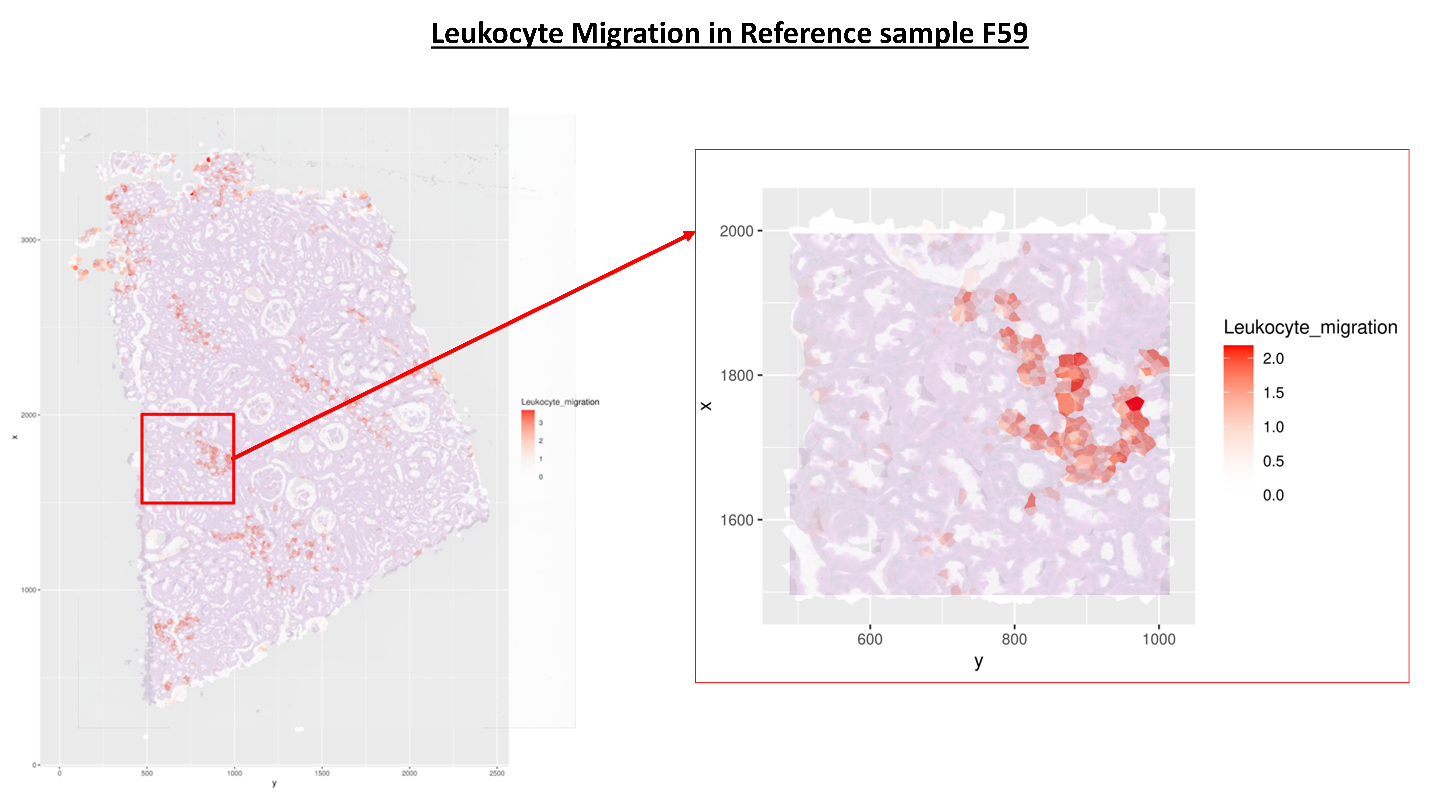

Supplementary Figure 20. Sample F59 – Reference, H&E image overlapped with pathway expression of leukocyte migration, showing both the full sample and a zoomed-in area of high pathway expression.

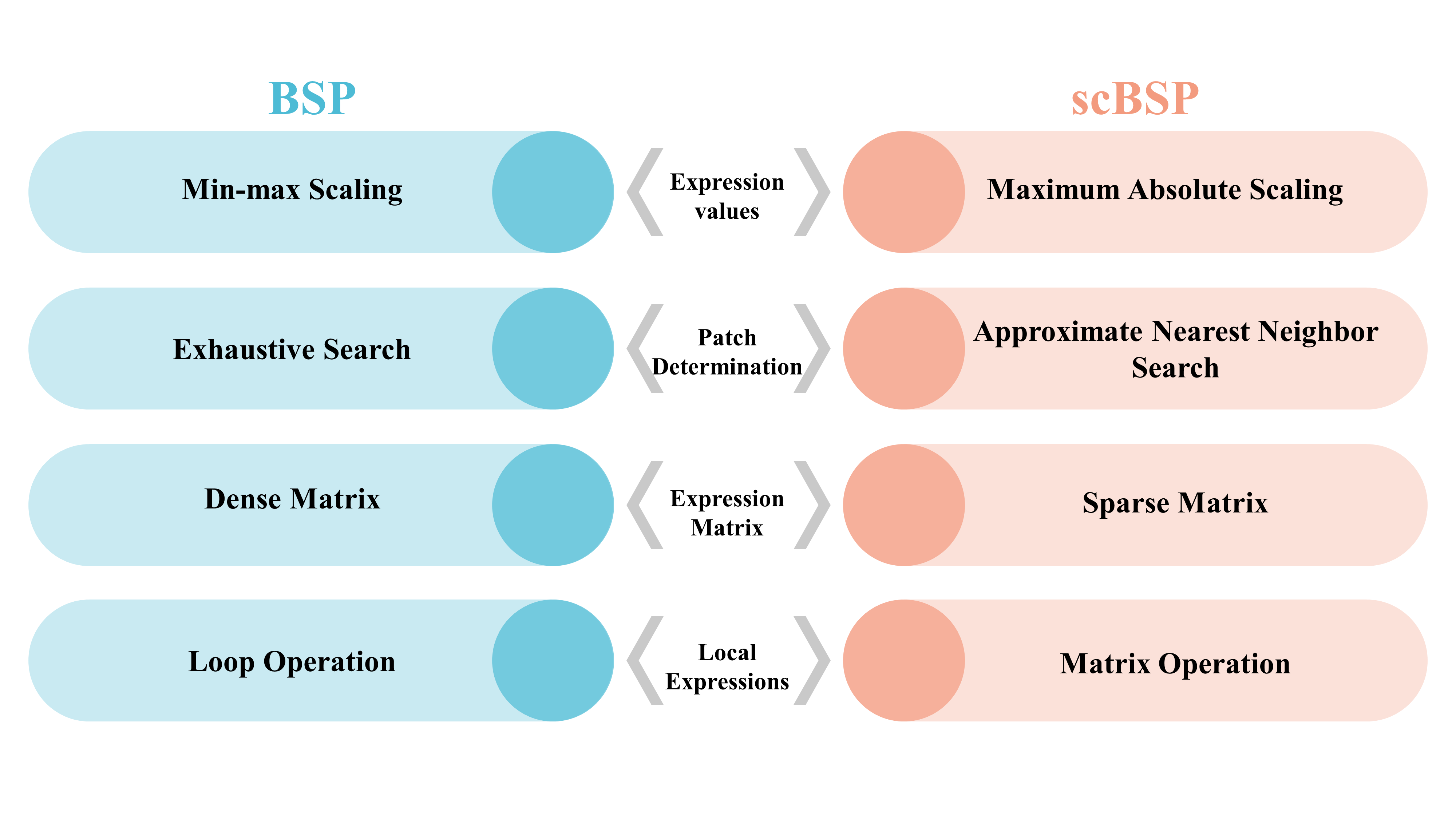

Supplementary Figure 21. Comparisons between scBSP and BSP.
